# The AccelerAge framework: A new statistical approach to predict biological age based on time-to-event data

**DOI:** 10.1101/2023.11.13.566803

**Authors:** Marije Sluiskes, Jelle Goeman, Marian Beekman, Eline Slagboom, Erik van den Akker, Hein Putter, Mar Rodríguez-Girondo

## Abstract

Aging is a multifaceted and intricate physiological process characterized by a gradual decline in functional capacity, leading to increased susceptibility to diseases and mortality. While chronological age serves as a strong risk factor for age-related health conditions, considerable heterogeneity exists in the aging trajectories of individuals, suggesting that biological age may provide a more nuanced understanding of the aging process. However, the concept of biological age lacks a clear operationalization, leading to the development of various biological age predictors without a solid statistical foundation. This paper addresses these limitations by proposing a comprehensive operationalization of biological age, introducing the “AccelerAge” framework for predicting biological age, and introducing previously underutilized evaluation measures for assessing the performance of biological age predictors. The AccelerAge framework, based on Accelerated Failure Time (AFT) models, directly models the effect of candidate predictors of aging on an individual’s survival time, aligning with the prevalent metaphor of aging as a clock. We compare predictors based on the AccelerAge framework to a predictor based on the GrimAge predictor, which is considered one of the best-performing biological age predictors, using simulated data as well as data from the UK Biobank and the Leiden Longevity Study. Our approach seeks to establish a robust statistical foundation for biological age clocks, enabling a more accurate and interpretable assessment of an individual’s aging status.

## 1 Introduction

Aging is a complex and multifaceted physiological process characterized by a gradual functional decline, leading to an increased risk of disease and mortality [1]. Although chronological age is a strong risk factor for aging-related diseases and mortality [2], there is high variation in the timing of disease onset and death in older individuals: some already experience a strong decline of their functional capability in their sixties and die soon thereafter, others remain healthy until their late nineties [3, 4]. This suggests that the rate at which we age varies between individuals and cannot be captured by chronological age alone. As a consequence, it has been postulated that the true underlying aging status of an individual can be captured by their ‘biological age’. This biological age is meant to be a holistic measure of aging: its value should not be driven by specific aging-related diseases, but should instead be a measure of one’s overall position in their total lifespan [5].

Individual aging, which is what biological age aims to capture, is often described using the metaphor of a clock: as if each of us possesses some latent clock that is ticking slowly but surely towards death, with a rate that varies between individuals. This clock paradigm is not new: Alex Comfort, widely accredited with being one of the founding fathers of the biological age prediction field [6], already used the term ‘age clock’ in 1969 [7]. Apparently, clocks are in line with the conceptual framework that we use to think about aging. In many recent publications, the terms ‘(biological) age predictors’ and ‘aging clocks’ are used interchangeably [8].

The quest for a reliable and valid biological age predictor is decades old [7, 9, 10]. Whereas the earliest attempts mainly used low-dimensional clinical variables as predictor variables, also known as ‘candidate markers’ (of aging), in the last decade the attention has started to shift towards high-dimensional candidate markers. This shift was initiated by the publication of Hannum’s and Horvath’s epigenetic age predictors [11, 12], which used DNA methylation (DNAm) data as the predictor variables and chronological age as the outcome. The difference between predicted age and true chronological age was found to be associated with various age-related outcomes and hence interpreted as a measure of biological aging [13]. Since then, various other chronological age-trained biological age predictors have been developed using different sources of omics [14–18].

As chronological age-trained approaches rely on a strong and untestable assumption [19], a second generation of biological age clocks have been developed, which use time-to-mortality as the outcome of interest. These mortality-trained predictors were found to outperform first-generation chronological age-trained predictors in their association with time-to-mortality, a broad range of common health conditions, physical and cognitive performance, age-related clinical phenotypes and frailty measures [20–23]. Two well-known second-generation predictors, both developed using DNAm data as candidate markers of aging, are PhenoAge [24] and GrimAge [25]. PhenoAge is a DNAm-based predictor of one’s ‘phenotypic age’, a constructed composite of nine clinical measures associated with time-to-mortality and chronological age. GrimAge is constructed by regressing time-to-mortality on a set of twelve DNAm-based surrogate markers for plasma proteins and smoking pack-years. The linear predictor of this model is linearly transformed such that the resulting predictions are on an age-scale.

Even though these second-generation clocks are an improvement over first-generation clocks, from a methodological point of view, both first- and second-generation clocks lack statistical rigour. When deciding on a prediction approach, it is important to first decide on the estimand. An estimand is the measure or quantity of interest that a predictor should predict. The concept of an estimand is important because it guides the choice of statistical methods and techniques to use. Clearly defining what measurable variable should be predicted is particularly important in the context of biological age prediction, as biological age is a latent concept, i.e. it is not directly observable. If this concept is not precisely defined, different people can therefore express different things with the term ‘biological age’. Therefore, the starting point of any biological age prediction approach should be to operationalize this latent concept into measurable variables or indicators: the estimands.

The currently existing biological age predictors, either age-trained or mortality-trained, did not follow from a solid operationalization of the concept of biological age. Hence, the estimand—the measurable quantity of interest that is to be predicted—is not clear. All biological age predictors claim to capture (a facet of) biological age, but if this concept is not properly operationalized it cannot be checked to what extent these predictors indeed predict what they (cl)aim to predict. In essence, these predictors are based on nothing more but a sequence of ad hoc computational steps. One consequence of this ad hoc nature is that these predictors are constructed using statistical models that are not in line with the conceptual framework of aging-as-a-clock that is so ubiquitous within the aging field. Another consequence of this missing operationalization is that biological age predictors cannot be properly evaluated: if it is unclear what should be captured by a predictor, it is not possible to formally check to what extent it is captured. At the moment, biological age predictors are generally only evaluated and compared by investigating to what extent the chronological age-independent part of the prediction (usually denoted by the symbol Δ and defined as the residuals after regressing predicted biological age on chronological age) is associated with time-to-mortality and other aging-related outcomes such as common health conditions, physical and cognitive performance, age-related clinical phenotypes and frailty [20–23]. Although this is a necessary condition for a biological age predictor, it is not a sufficient one: biological age predictions should not only be meaningful with regards to some other variable (as is the case when only evaluating Δ), but these predictions themselves should also be directly interpretable. To be able to compare the performance of biological age predictors on this characteristic, additional evaluation measures need to be considered.

Given the gaps and limitations of current biological age predictors, namely that they are not based on a proper operationalization of biological age, are not in line with the aging-as-a-clock paradigm and are difficult to properly evaluate, the aim of this paper is threefold. Firstly, we propose an operationalization of biological age. Secondly, we present the *AccelerAge* framework: an new approach to predict biological age that follows directly from the proposed operationalization of biological age. This provides AccelerAge with a solid statistical foundation and makes it more than just a sequence of computational steps. The AccelerAge framework is based on Accelerated Failure Time (AFT) models [26]. Unlike current second-generation prediction approaches, which rely on Proportional Hazard (PH) models [27], AFT models model the effect of candidate markers of aging on one’s survival time directly. As we will illustrate, this assumed accelerating or decelerating effect of candidate markers on survival is directly in line with the clock metaphor that is so ubiquitous in aging research. The appeal— but underuse—of AFT models in the context of aging research has been noted before [28], but not in the context of biological age prediction. Thirdly, we introduce two new evaluation measures for biological age predictions. These evaluation measures consider the association between the age-independent part of biological age predictions and mortality and directly evaluate the biological age predictions themselves. Finally, for illustration purposes we will build predictors based on our AccelerAge framework and compare these to a predictor built by taking a similar approach as was taken to build GrimAge, which is currently considered the best-performing biological age predictor. The performance of the predictors based on the AccelerAge framework and on GrimAge are evaluated and compared using simulated data as well as data from the UK Biobank and the Leiden Longevity Study. With this we hope to contribute to a solid statistical foundation underpinning biological age clocks.

## 2 Biological age operationalization

Biological age should be a measure of one’s position in their total life or health span [5]. One of the earliest papers discussing biological age, from Benjamin [9], already defined biological age as such: “Biologic age may be defined once more through an example: A man (white, American, of upper cultural level) who has lived seventy years has a life expectancy of about nine years. He has, therefore, a chance to celebrate his seventy-ninth birthday. These are statistical figures based on national mortality reports of many thousands of cases. The individual is not considered except as part of the population. The examination of this man shows that he has a favorable heredity, sound organs, and that he functions like a much younger man. The age estimation (to be described later on) makes him 15 years younger, that is to say, his condition corresponds to that of a man of 55, which would be his approximate biologic age.”

This idea to define biological age through life expectancy and a comparison with some reference population is intuitively appealing. However, a formal operationalization is required, such that a proper prediction approach can be decided upon, resulting in a predictor that predicts this operationalized concept. Our proposal for a formal mathematical definition of biological age is given below.

Denote by *B* biological age, by *C* chronological age, by *X* a (set of) true marker(s) of biological age (defined as markers that, conditional on *C*, are associated with *B*) and by *T* age-at-death. Under our proposed operationalization, *B* is defined through some function of residual life. We consider mean residual life (mrl): define mean residual life at age *t* as mrl(*t*) = *E*(*T − t|T > t*). Now we can define biological age as follows: individual *i*, with chronological age *C* = *c_i_*and marker value *X* = *x_i_*, has *B* = *b_i_* if

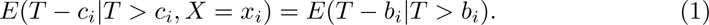

This can also be written as: *B* = *b_i_*if

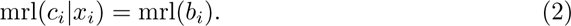

Hence, we assume that biological age is closely related to expected residual life, and is defined with respect to some reference population: given a prediction of mrl(*c_i_|x_i_*) and a population lifetable of some reference population, biological age is determined by checking which chronological age within the population corresponds to a mean residual life value of mrl(*c_i_|x_i_*). For example, a heavy smoker aged 50 might have a life expectancy of 20 years given his marker values *X_i_*, while in the general male population a life expectancy of 20 years corresponds with an age of 57. Then the heavy smoker’s biological age is defined as 57. This is visualized in Figure 1.

**Fig. 1.**
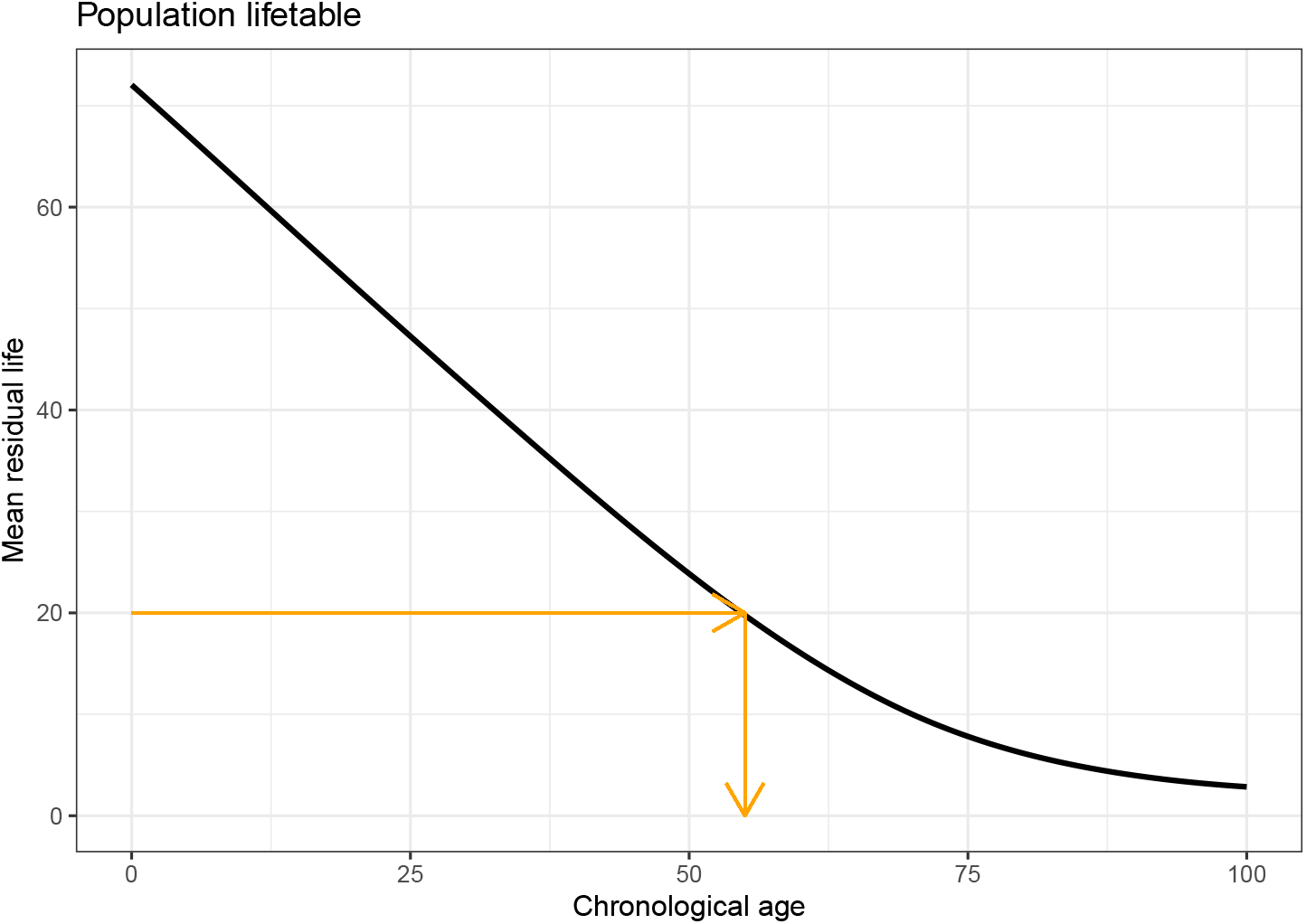
Illustration of step 2 of our biological age operationalization: going from a mean residual life prediction to a biological age prediction. The black line denotes mean residual life at chronological age *t* within some reference population. Someone with an estimated mean residual life of 20 years has a corresponding biological age of 57.

This operationalization hence suggests a two-step approach for prediction of *B*: the first step is to obtain 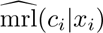 by regression-based estimation of mrl(*t|X*); the second step is to obtain 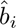 using mrl(*t*), which is generally taken from some external source.

Under this operationalization, someone’s biological age is per definition determined with respect to some reference population. This allows for a meaningful and direct interpretation of predictions: a biological age of 50 means that someone has “the same life expectancy as the average person with chronological age 50 *within the reference population*”. If the reference population changes, someone’s mean residual life prediction will not change, but the resulting biological age prediction will. A possible disadvantage of this is the flip side of the same coin: choosing a reference population might not always be straightforward and/or the appropriate lifetable might not always be available. However, this can be circumvented by using the training data set itself as a reference population, if it is large enough and covers a wide enough chronological age range.

For convenience’ sake we have considered mean residual life (mrl) here, but one could also use some other function of residual life, e.g. median residual life (medrl), without loss of generality. When considering time-to-mortality as the outcome of interest, we have found the differences between mrl and medrl to be negligible. The choice for a particular function of residual life can be based on practical arguments: e.g. availability of a corresponding lifetable (most lifetables provide mrl) and computational speed (medrl is faster to compute).

There exist several statistical models that can be used to obtain conditional residual life estimates 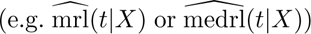 using time-to-event data. In this paper we will do so using Accelerated Failure Time models, using chronological age as the timescale *t*, which we believe to be a natural fit to the ‘aging as a clock’ paradigm. In contrast, for the GrimAge predictor the more frequently used semi-parametric Cox Proportional Hazards model is used, using time-on-study as the timescale *t*. We describe both models in more detail in the next section.

## 3 Proportional Hazards and Accelerated Failure Time

When working with survival data, two commonly encountered regression models are Proportional Hazards (PH) and Accelerated Failure Time (AFT) models. They differ in the assumption on how predictor variables act on one’s survival. In this section we describe both models in detail.

Proportional Hazard (PH) models, due to Cox [27], are the most commonly used models for survival analysis. In this model, survival is modeled through the hazard function *h*(*t*), also known as the instantaneous failure rate. PH-based models assume that the exponent of the linear predictors has a multiplicative effect on the hazard:

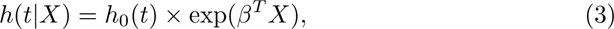

where *h*_0_(*t*) denotes the baseline hazard. This implies the effect of the linear predictors on the survival curve *S*(*t*) is given by:

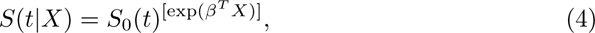

where *S*_0_(*t*) denotes the baseline survival. The semi-parametric Cox PH model is the most frequently encountered PH model; it is semi-parametric because it does not make assumptions about the specific form of the baseline hazard function, only about the effect of covariates on this hazard.

AFT models provide an alternative to PH models for the modelling of survival data [26]. In contrast to PH models, the AFT approach models survival times directly. AFT models assume that the effect of covariates on the baseline survival curve is to shrink or stretch this curve, i.e. accelerate or decelerate it by a factor exp(*β^T^ X*):

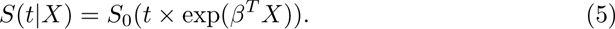

In contrast to PH models, which are often semiparametric (i.e. no parametric distribution is assumed for the baseline hazard *h*_0_(*t*)), it is common to fit parametric AFT models, which assume that the survival times *T* follow a known parametric distribution. In principle parametric models are more restrictive than semi-parametric models. However, for models where the event of interest is known to follow a certain distribution, it can be an advantage. For adult mortality this is the case: it has long been known that adult mortality (of many different species, amongst which humankind) is accurately described by the Gompertz distribution [29], with baseline hazard function *h*_0_(*t*) = *a* exp(*bt*). A lack of fit has been reported for the extreme old, known as ‘late-life mortality deceleration’—this phenomenon was in fact already reported by Gompertz himself [30]—but others have found this deceleration to be negligible until over 100 years of age [31]. A more recent study concluded that this mortality deceleration is notoriously difficult to prove [32]. Although it is sometimes stated that the Gompertz distribution can only be parameterized as a PH model, this is not correct: it can also be parameterized as an AFT model [33]. This is illustrated in section 1 of the Supplementary Information.

When modelling survival data, a choice has to be made about the appropriate timescale *t*. If subjects are followed from some well-defined event, e.g. randomization in a clinical trial, the relevant timescale is time-on-study and all subjects enter at time *t_start_* = 0. In the context of cohort studies, however, it has long been argued that chronological age is the preferred timescale [34, 35]. Nevertheless, the PH model used in the construction of the GrimAge predictor uses time-on-study at the timescale *t* (we refer section 2 of the Supplementary Information for more details).

The appeal of AFT models in the context of biological age prediction is that the factor exp(*β^T^ X*) in Equation (5) can be directly interpreted as an individual aging rate. If exp(*β^T^ X*) is greater than 1, a subject experiences accelerated aging: time *t* is multiplied by exp(*β^T^ X*). If exp(*β^T^ X*) is smaller than 1, a subject experiences decelerated aging. AFT models therefore tie in nicely with the clock paradigm, as they provide an intuitive measure of accelerated aging (a faster ticking clock) or decelerated aging (a slower ticking clock).

In Figure 2 we visualize the effect of covariates on the hazard rate and the survival curve for both AFT and PH models. In the top row, the black line represents the baseline survival curve. In the bottom row, the black shade represents the corresponding baseline hazard. The artificial plateau in midlife (where no one dies, i.e. the hazard is zero and hence the survival curve remains constant) was added to more clearly illustrate the difference between AFT and PH models. The topleft panel shows the effect on the survival curve for someone with ‘beneficial’ covariates under the PH model: i.e. exp(*β^T^ X*) is negative. At every point in time, this person experiences a lower hazard, because the baseline hazard is multiplied by a factor exp(*β^T^ X*), in line with Equation (3). Hence, at every given age baseline survival is shifted upwards, because it is raised to the power *−* exp(*β^T^ X*): it is as if this person is protected by some shield. The location of the plateau in midlife, however, does not change. The topright panel shows the effect on survival for someone with beneficial covariates under the AFT model. Now the survival curve is stretched out in the horizontal direction, in line with Equation (5): the biological age clock of this person is ticking a factor exp(*β^T^ X*) slower than the clock of the baseline population. As a result this person also experiences the hazard-free period in midlife at a later chronological age. The AFT model is hence a more natural fit to the aging-as-a-clock concept than the PH model.

**Fig. 2.**
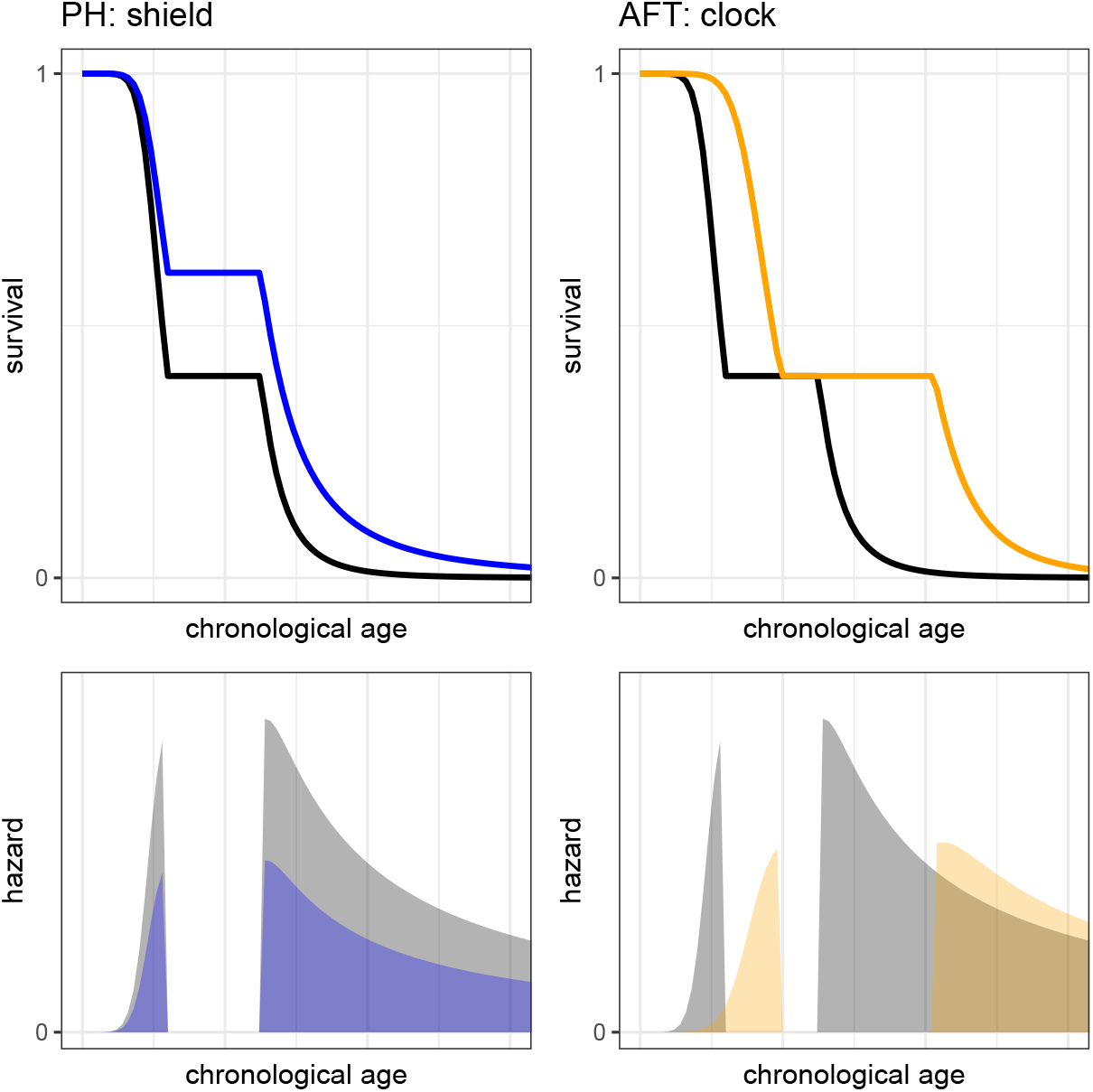
Illustration of the effect of markers on baseline hazard and survival under the assumption of Proportional Hazards (left panels) and Accelerated Failure Time (right panels).

## 4 The AccelerAge framework

In this section we introduce our new statistical framework for biological age prediction, using an Accelerate Failure Time model with chronological age as the timescale *t*. This framework is in line with our suggested operationalization for biological age as given in section Biological age operationalization: i.e. biological age is based on (mean) residual life and determined relative to some reference population. This suggests a two-step approach for prediction of *B*:

1. Get 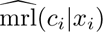 by regression-based estimation of mrl(*t|X*);
2. Obtain 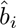 using mrl(*t*) (generally available from some external source) so that 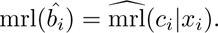

The first step in arriving at a prediction for biological age is to define a model for mean residual life mrl(*t|X*), given that a subject has survived until time *t*. Here, *X* denotes a (set of) true marker(s) of aging. We choose an approach via the survival function. In terms of the survival function *S*(*t*), mrl(*t|X*) can be expressed as:

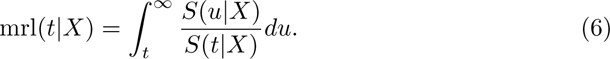

How exactly *X* affects *S*(*t|X*) depends on the underlying model. Since we assume the AFT assumption to hold, this relationship is given by Equation (5).

For step 2, one must have access to the population lifetable of some reference population, in order to translate the mean residual life prediction to a biological age prediction. We suggest to use the lifetables as provided by the national statistics office of the country where the data was gathered. Alternatively, in some cases the data set used to fit the model in step 1 could also be used to construct a life table.

We call this approach the AccelerAge framework, to emphasize its close link with Accelerated Failure Time models. We chose the term ‘framework’–in the sense of a conceptual structure–to emphasize the fact that ‘AccelerAge’ refers both to the operationalization of biological age as well as to the modelling approach that is followed to construct a predictor. Hence, we distinguish between a *framework* (the structure used to define biological age), a prediction *approach* (all modelling steps required to arrive at a prediction) and a *predictor* (a fitted version of a particular framework, fitted on a particular dataset, following all steps of the modelling approach).

In this paper we deliberately present a framework instead of a predictor: our AccelerAge framework can be applied to any type of time-to-event data using any kind of predictor variables. The AccelerAge framework is based on a proper operationalization of biological age, is in line with the ‘aging as a clock’ paradigm and its predictions are directly interpretable on an age-scale. The names ‘GrimAge’ and ‘PhenoAge’ are generally considered to refer to predictors: these names refer to a specific fitted model, which can be applied to a specific set of predictor variables (DNA methylation data) to arrive at a prediction. Nevertheless, in principle the modelling approaches that were followed to construct GrimAge and PhenoAge can also be used in different settings (e.g. with different predictor variables) to produce ‘GrimAge-type’ and ‘PhenoAge-type’ predictors.

## 5 Evaluation measures

The standard approach in the evaluation of biological age clocks is to check whether the chronological age-independent part of a biological age prediction 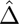 (also known as the ‘age acceleration’ and obtained by regressing 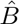 on *C*) is associated with mortality and other aging-related outcomes, such as frailty or cardiovascular disease. However, the biological age prediction 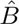 should not only be meaningful in relation to some other time, which is what is done when evaluating 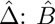 itself should also be meaningful. In this section we therefore introduce two new evaluation measures of biological age predictions which evaluate this: discrimination and calibration. Both are routinely evaluated aspects of (clinical) prediction models based on survival data [36].

To check to what extent individuals with a higher predicted biological age indeed die sooner—in other words, to what extent the predictor is able to discriminate—we propose to consider the concordance (also called C-index) of a predictor. The concordance can be interpreted as the fraction of pairs in the data where the observation with the higher observed residual life also has the lower biological age. We use Uno’s C-index, which does not depend on the study-specific censoring distribution [37].

To check to which extent the biological age predictions are on the proper scale— in other words, to what extent the predictor is well-calibrated—we propose to use calibration plots. For any biological age prediction, it is possible to obtain the corresponding *X*-year mortality probability for this age using the population lifetable. Individuals can then be grouped based on their predicted *X*-year mortality probability in *N* equally sized groups. Per group, the mean predicted mortality probability can be compared with the true observed mortality rate within this group. If the predictor is well-calibrated, these correspond closely.

## 6 Simulation study set-up

We conducted a simulation study to check the predictive performance of predictors fitted using the AccelerAge framework and compared it to a predictor fitted using the same approach as was used to construct GrimAge. GrimAge is currently considered the best-performing biological age predictor, as the age-independent part of GrimAge predictions has been found to be associated with more aging-related outcomes than the age-independent part of PhenoAge-predictons [20, 22].

We generated data under a similar study design as that of real data sets used to train longitudinal biological age clocks, namely prospective cohort studies: people enter the study at a random chronological age *C* and are then followed-up over time. For each individual in our simulated data set we generate their marker values *X*, their age-of-death *T* (the distribution of which depends on the chosen baseline) and the chronological age *C* at which we would include them in our study. By excluding individuals for which *T < C*, we mimic the selection process that also takes place in real prospective cohort studies: people who have already died cannot be observed.

Data was generated assuming two true markers *X*_1_ and *X*_2_, constant over each person’s lifetime and following a standard normal distribution at birth. Although the assumption that markers are constant over one’s lifetime is not a realistic one (most omics-based markers change over the course of a lifetime), it is in line with the original approach followed to construct the GrimAge predictor, in which the the composite markers are age-adjusted before inclusion in the PH model.

We considered three data-generating mechanisms: one in which the baseline hazard *h*_0_(*t*) follows a Gompertz distribution and the PH assumption holds (referred to as Gompertz-PH), one in which *h*_0_(*t*) follows a Gompertz distribution and the AFT assumption holds (referred to as Gompertz-AFT) and one in which *h*_0_(*t*) follows a Weibull distribution (referred to as Weibull), which per definition can be parameterized as both an AFT and a PH model at the same time. We included the Gompertz scenarios because the Gompertz distribution is known to accurately describe (human) lifespan. We included the Weibull scenario to illustrate the effect of a misspecified baseline when fitting a parametric AFT model.

We independently generated observations of markers *X*_1_ and *X*_2_ and chronological age *C* as follows:

- 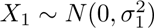 — biomarker, constant over one’s lifetime;
- 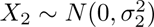 — biomarker, constant over one’s lifetime;
- *C ∼ U* (*c_min_, c_max_*) — chronological age at which individual would enter the study.

We used the following parameter values: *σ*_1_ = *σ*_2_ = 1, *c_min_* = 20 and *c_min_* = 80. Next we generated age-at-death *T* under each of the three data-generating mechanisms. For a given individual *i*, under the PH-assumption, age-at-death *t_i_* given *X_i_* (where *X_i_* here denotes a vector of *x_i_*_1_ and *x_i_*_2_) can be drawn as follows:

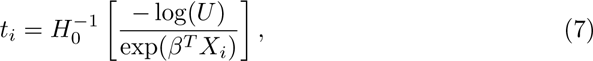

where *U* follows a uniform distribution on the interval from 0 to 1. Under the AFT-assumption, *t_i_* given *X_i_* can be generated as follows:

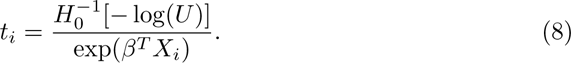

We provide the derivation of Equations (7) and (8) in section 3 of the Supplementary Information. The expression for 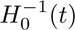 depends on the chosen baseline distribution: Gompertz for the first two scenarios, Weibull for the third. We chose the parameters of these distributions such that the resulting event times approximately resembled human lifespan: for Gompertz, *a* = exp(*−*9) and *b* = 0.085 (where the baseline hazard is given by *h*_0_(*t*) = *a* exp(*bt*)), for Weibull, *λ* = 34*^−^*^10^ and *ν* = 8 (using the operationalization as given in Bender et al. [38], where *h*_0_(*t*) = *λνt^ν−^*^1^). For Gompertz-PH, *β* = (0.3, 0.3), for Gompertz-AFT, *β* = (0.05, 0.05) and for Weibull, *β* = (0.35, 0.35). These coefficients cannot be directly compared between the three scenarios, since on a PH-scale the interpretation of an effect size is different than on an AFT-scale, but they were chosen such that the resulting age-of-death distribution was comparable between the three scenarios, as can be seen in Figure 3

**Fig. 3.**
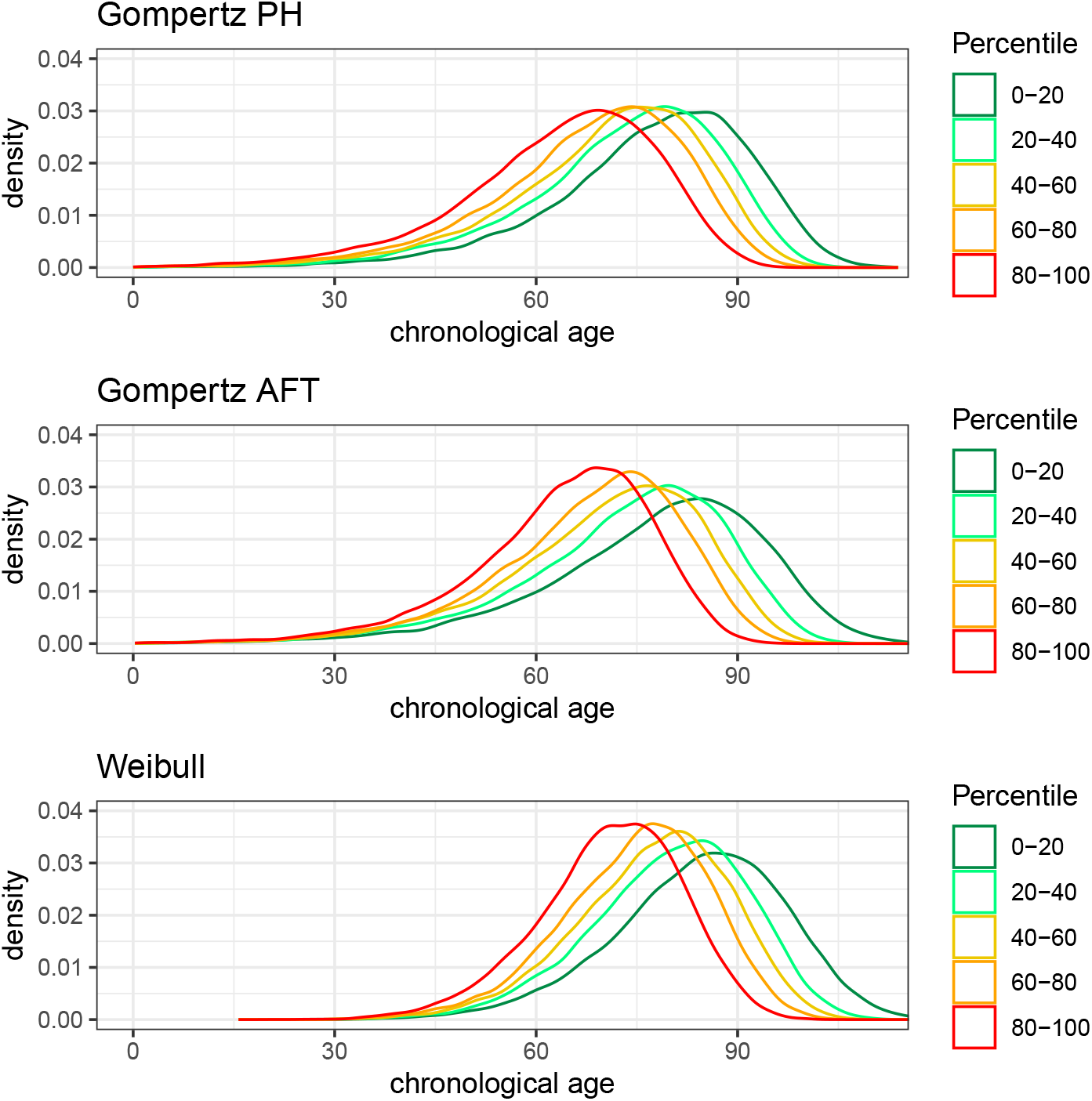
Histogram of the ages-of-death (generated at birth) for different quantiles of the linear predictor for each of the three data-generating mechanisms considered.

If for a given individual age-at-death *t_i_* was smaller than chronological age *c_i_*, this individual is not observed, since he or she has already died at the age that we otherwise would have observed them. Those cases were removed. For the remaining individuals, we determined their survival curve *S_i_*(*t*) via Equations (4) or (5).

We consider median residual life (medrl) instead of mean residual life (mrl) because medrl is considerably faster to compute than mrl, since there is no integration involved. A pilot simulation was conducted considering both mrl and medrl: results were very similar under the settings considered in this simulation. For each individual in the data set we determined 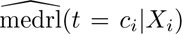 as follows. First, determine the time *t* = *t_i,med_* at which survival is half the current value *S_i_*(*t* = *c_i_*). Next, subtract *c_i_* from *t_i,med_* to obtain expected median residual life medrl(*c_i_|X_i_*).

The final step in simulating biological age *B* involves a population lifetable for median residual life, medrl(*t*). The lifetables were constructed using the true parameter values of the three different data-generating mechanisms. Finally, for each subject *i*, we determined their biological age by checking in the population lifetable for which *t* the median residual life prediction of a particular individual 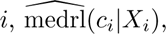 is equal to the population’s medrl(*t*).

We assumed individuals were followed-up for a period of 20 years. So if (*t_i_ − c_i_*) *>* 20, the age-of-death of this individual is censored and he/she is observed until age *c_i_* + 20. If (*t_i_ − c_i_*) *<* 20, this individual is observed until their age-of-death *t_i_*. Figure 4 contains plots of chronological age against biological age for a realization of each of the three data-generating mechanisms described.

**Fig. 4.**
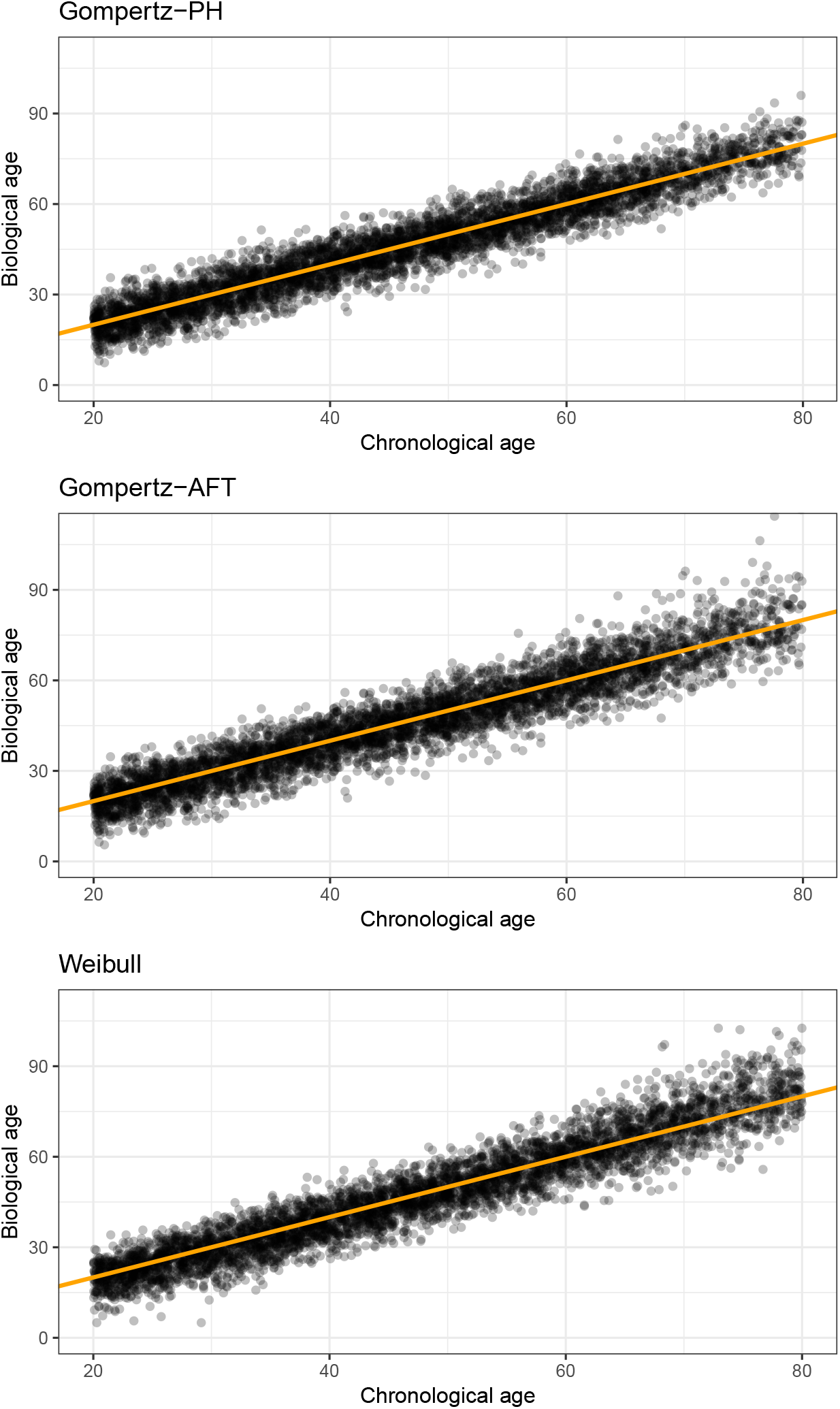
Plots of chronological age against (true) biological age for a simulated dataset of size *n_obs_* = 5000 for each of the three data-generating mechanisms considered.

We generated training data sets of sizes *n_obs_* = 500, 2500, 5000, 7500 and 10,000 under each of the three data-generating mechanisms (Gompertz-AFT, Gompert-PH and Weibull) and generated test sets of size *n_test_* = 5000.

Three different prediction approaches were used to fit three different biological age predictors on the simulated training datasets: 1) a predictor based on the AccelerAge framework with a Gompertz baseline (AccelerAge-Gompertz), 2) a predictor that, like the AccelerAge framework, works via residual life and lifetables but assumes proportional hazards (PH-semipar) and 3) a predictor constructed following the approach that was used to construct the original GrimAge predictor (GrimAge-type). An elaborate description of the approach we took to fit this predictor can be found in section 2 of the Supplementary Information. Note that of these three prediction approaches, GrimAge-type is the only one that uses time-on-study as timescale *t* and relies on an ad hoc transformation to an age scale.

AccelerAge-Gompertz is correctly specified for the Gompertz-AFT data-generation mechanism. PH-semipar is correctly specified for the Gompertz-PH data-generating mechanism. Since PH-semipar is semiparametric, it is also correctly specified for the Weibull data-generation mechanism. GrimAge-type is not based on an underlying definition or operationalization of biological age, so it is not clear under which data-generation mechanism it would be correctly specified. But since it uses a Cox PH regression, it is to be expected that it will do better under the Gompertz-PH and Weibull data-generating mechanisms than under the Gompertz-AFT data-generating mechanism.

We use root-mean-square error (RMSE) as the performance measure: 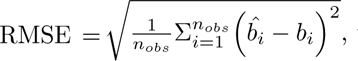 where for individual 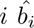 denotes predicted biological age and *b_i_* denotes true biological age. The simulation study sample size is *n_sim_* = 200 (i.e. 200 simulated data sets of size 500, 2500, 5000, 7500 and 10,000 for the three different data-generation mechanisms is 200 *×* 4 *×* 3 = 2400 simulated training data sets in total). In a few cases the Gompertz AFT model could not be fitted on the simulated data set, as it is a numerically delicate model to fit: an error was thrown that the Hessian was singular or that the model did not converge. This was the case for 9 of the 2400 simulated training data sets. We left those data sets out. For each data-generation mechanism and for each *n_obs_*-size we report the average RMSE over the *n_sim_*repetitions.

All analyses were performed using R version 4.1.0. The parametric AFT models were fitted using the eha package version 2.10.3 [39].

## 7 Simulation study results

The results for the Gompertz-PH, Gompertz-AFT and Weibull data-generating mechanisms are presented in Figures 5, 6 and 7, respectively.

**Fig. 5.**
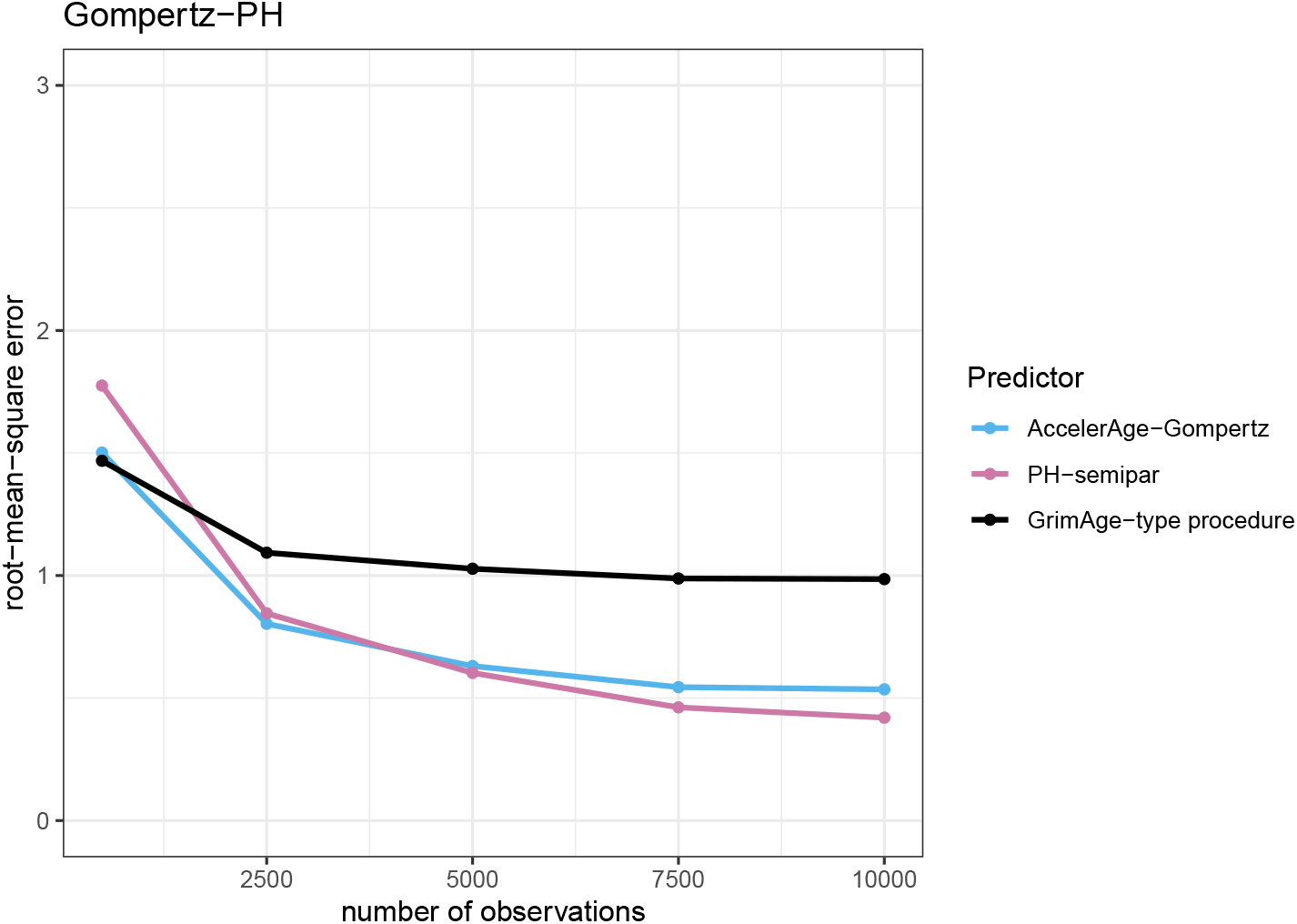
Performance of the three different biological age predictors in terms of the root-mean-square error under the Gompertz-PH data-generating mechanism. Results are reported for data sets of varying sizes (*n_obs_* = 500, 2500, 5000, 7500 and 10,000) as the average root-mean-square error over a simulation sample size of *n_sim_* = 200.

**Fig. 6.**
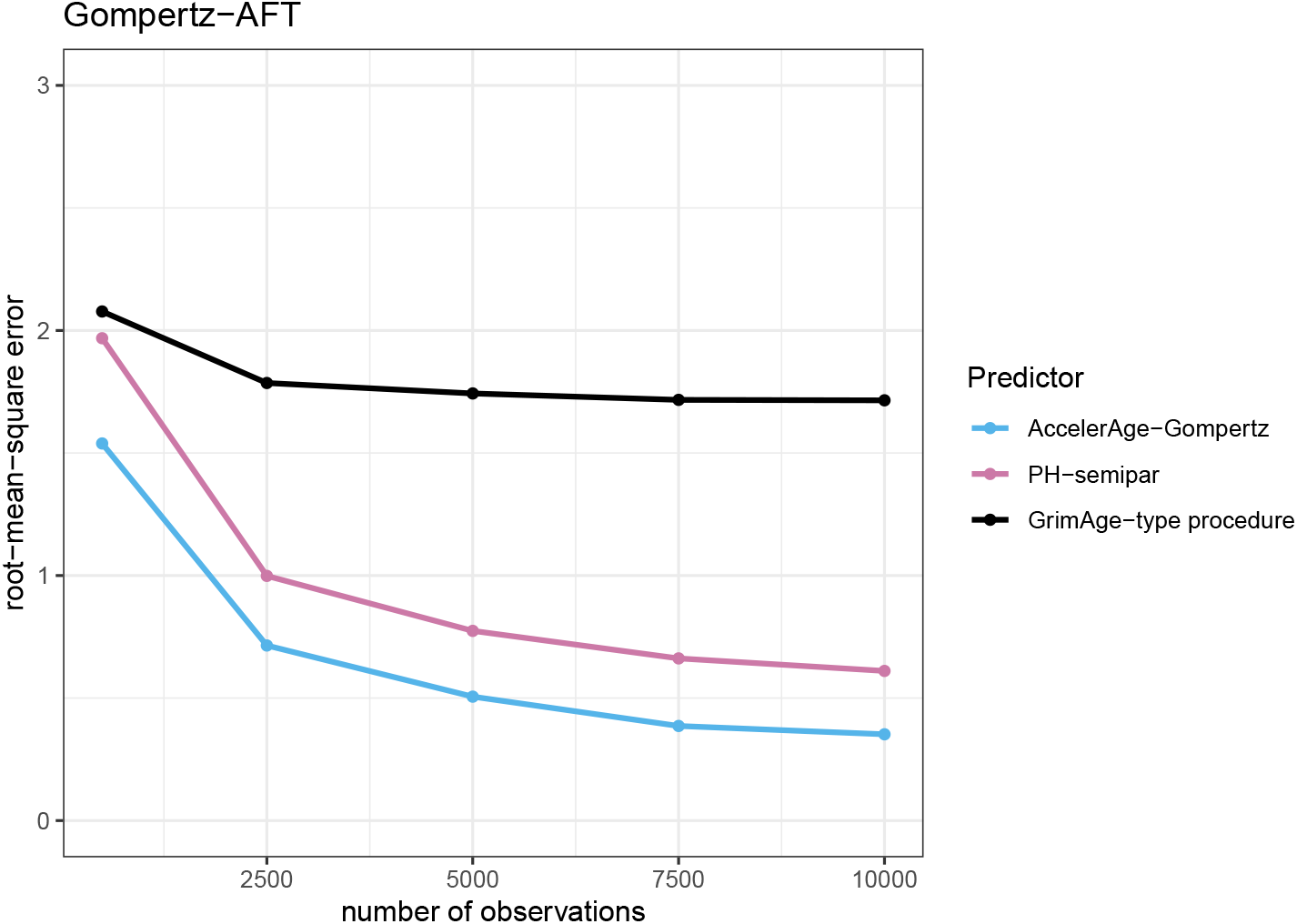
Performance of the three different biological age predictors in terms of the root-mean-square error under the Gompertz-AFT data-generating mechanism. Results are reported for data sets of varying sizes (*n_obs_* = 500, 2500, 5000, 7500 and 10,000) as the average root-mean-square error over a simulation sample size of *n_sim_* = 200.

**Fig. 7.**
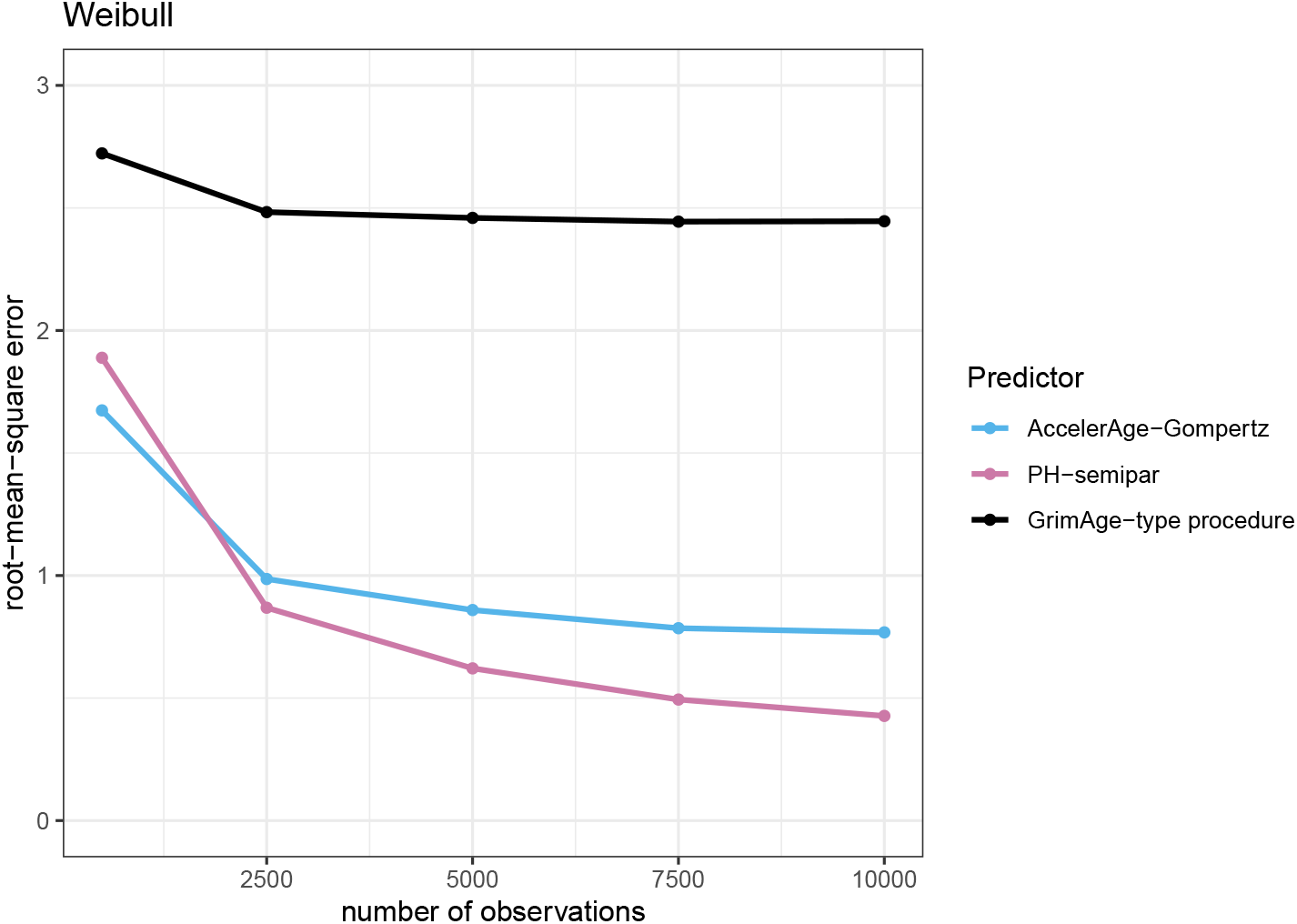
Performance of the three different biological age predictors in terms of the root-mean-square error under the Weibull data-generating mechanism. Results are reported for data sets of varying sizes (*n_obs_* = 500, 2500, 5000, 7500 and 10,000) as the average root-mean-square error over a simulation sample size of *n_sim_* = 200.

Figure 5 shows that under the Gompertz-PH data-generating mechanism, GrimAge-type does best if the training data sample size is small (*n_obs_*= 500): the RMSE is lowest. This is likely due to the fact that GrimAge-type uses a Cox PH model to estimate the effect of candidate markers on mortality, which matches the used data-generating mechanism here. GrimAge-type does not need to estimate the population baseline survival *S*_0_(*t*) to arrive at biological age predictions due to its ad hoc transformation of the linear predictors. The other predictors do: 500 observations in a training data set (of which a substantial number censored) apparently do not suffice to properly estimate baseline survival. This is advantageous to GrimAge-type. However, whereas the RMSE of the other predictors keep decreasing with increasing training dataset size,

GrimAge-type’s performance stops improving. AccelerAge-Gompertz and PH-semipar perform similarly and outperform GrimAge-type when the size of the training data is larger than approximately 1,500 samples. Although PH-semipar is correctly specified under this data-generating mechanism, this predictor also needs enough events to properly estimate *S*_0_(*t*). And indeed, as *n_obs_* increases, PH-semipar eventually outperforms AccelerAge-Gompertz. Even though AccelerAge-Gompertz is misspecified under this data-generating mechanism, its performance is still quite good. This can likely be attributed to the fact that it assumes the correct underlying baseline distribution and the effect sizes *β* are relatively small.

In Figure 6 it can be seen that under the Gompertz-AFT data-generating mechanism, the corresponding correctly specified predictor (AccelerAge-Gompertz) performs best from the start. GrimAge-type again does okay for smaller training data sets but here its performance also quickly stops increasing.

Figure 7 shows that under the Weibull data-generating mechanism, GrimAge-type performs worst of all predictors with a large margin. This is likely due to the fact that the Cox PH regression in GrimAge’s second step uses time-on-study as the timescale instead of chronological age. This affects its performance. This mismodeling was less of an issue for the Gompertz-PH and Gompertz-AFT scenarios in Figures 5 and 6, because the Gompertz distribution belongs to the so-called ‘exponential family’. For this family mismodeling of time in a Cox PH model does not matter (for an elaborate discussion, see Thíebaut and Bénichou [35]). The performance of the AccelerAge predictor and PH-semipar is in line with expectations: for larger data sets, the correctly specified PH-semipar performs best, closely followed by AccelerAge-Gompertz.

## 8 Real data illustration

In this section we evaluate and compare the performance of a predictor fitted using our newly proposed AccelerAge framework and a predictor fitted using the GrimAge framework on real data. We use data from the UK Biobank (UKB). The UKB is a large population-based prospective cohort study. Between 2006 and 2010, more than 500,000 participants aged 37–73 were recruited from different sites across the United Kingdom. Participants’ health is being followed long-term. The study contains extensive phenotypic and genotypic detail about its participants, including biological measurements and biomarker values, and longitudinal follow-up for a wide range of health-related outcomes, provided through a linkage with medical and health records[40].

We use blood-based metabolic biomarkers as predictor variables. The metabolic biomarkers were quantified using the high-throughput nuclear magnetic resonance (NMR) targeted metabolomics platform of Nightingale Health Ltd. (Helsinki, Finland), known for its high repeatability over time and absence of batch effects [41, 42] Per EDTA plasma sample, 249 metabolic measures are provided (of which 168 in absolute concentration units and 81 ratios). The majority of these biomarkers relate to lipoprotein metabolism. In the UKB the first tranche of NMR-metabolomics data became available in March 2021, for more than 120,000 samples (from approximately 118,000 participants at baseline and 4,000 at repeat assessment, of which 1,500 at both) [43]. We only included those subjects with measurements at baseline. Metabolic variables for which measurements were missing for more than 1 percent of all samples were excluded (excluding 7 of 249). Subjects with missing measurements in the remaining 242 metabolic variables were also excluded (excluding 1,648 of 104,312). This left a sample of size N = 102,664, of which 47,300 men and 55,364 women. Mean age at recruitment was 56.3 years (sd = 8.1).

The absolute concentrations of the metabolic variables measured at baseline are known to be 5-10% diluted in the UKB data [44]. To still allow for validation of our fitted biological age predictors in an external dataset, we decided to scale and center each metabolic variable prior to analysis.

Prospective mortality (i.e. time-to-mortality) is the outcome of interest. Followup data until November 2021 was available. Median follow-up time was 13.3 years (IQR: 12.5–14.0). During follow-up 7,629 participants died. No participant was followed for more than 16.9 years. Since at inclusion no participant was older than 69, the population survival curve of the participants is only defined from age 40 to age 85. That the population survival curve is not well-defined at its right tail has significant negative consequences for the semiparametric PH-semipar predictor considered in the simulation study. As there is no information in the data on how the right tail of the survival function looks like in reality, these predictors cannot properly estimate baseline survival *S*_0_(*t*). We therefore decided to exclude PH-semipar from this real data illustration. This leaves two predictors: one based on our our AccelerAge Gompertz framework fitted on the metabolic biomarkers (referred to as metabo-Accelerage-Gompertz) and one based on the GrimAge approach, as described in the previous section, but now fitted on the metabolic biomarkers (referred to as metabo-GrimAge).

To construct population lifetables, necessary for the residual-life based biological age prediction approaches, we used data from the United Kingdom’s Office for National Statistics [45]. When comparing survival in the UKB population with that of the general population, it becomes apparent that the UKB participants on average live longer. This is shown in Figure 8, using the most recent period lifetables (2018-2020), stratified by sex, from the Office for National Statistics as a comparison. We scaled these curves such that it only starts decreasing from age 40 onward, to avoid an unfair comparison due to the immortal time bias present in the UKB data, as no participant was included before age 40. This figure also illustrates the limited age range of the UKB participants’ survival curves. The fact that UKB participants are not representative of the general population, but seem to belong to a healthier subset, has been noted before; see e.g. Fry et al. [46].

**Fig. 8.**
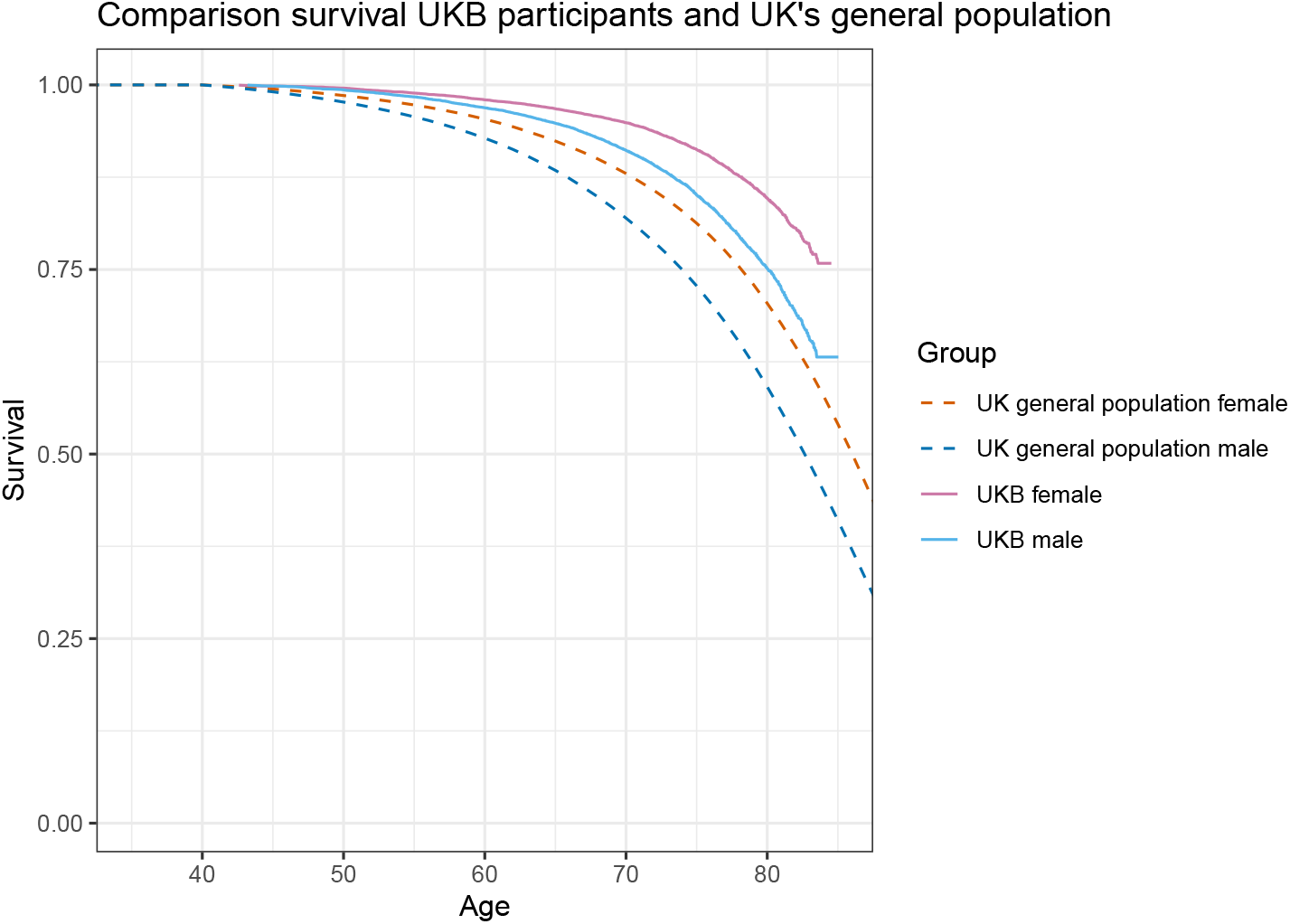
Comparison of survival curves of the included UKB participants, stratified by sex, and of the UK general population (the period lifetable of 2018-2020), stratified by sex, as provided by the Office for National Statistics.

The fact that we use sex-stratified population survival curves means that a man and a woman who have the same predicted mean residual life, will have a different predicted biological age: women live longer, so the woman’s biological age will be higher.

Given the large sample size available, we randomly selected 60,000 individuals to serve as the training data set and the remaining 42,664 individuals to serve as the test data set. As the metabolic variables are often strongly correlated, we fit our predictors on the training data using as predictor variables the first 22 principal components of the metabolic variables (together explaining at least 95 percent of the variance) and sex. Time-to-all-cause-mortality is the outcome.

We evaluated the biological age clocks using both standard and new evaluation measures, as introduced in section Evaluation measures. To calculate the concordance, we translated predicted biological age back to predicted mean residual life for metabo-AccelerAge-Gompertz, via the sex-specific baseline survival curves. For metabo-GrimAge it is unclear whether predictions can be directly translated to a predicted mean/median residual life value. We did try this, but it resulted in a lower concordance than when using the predicted GrimAges directly. We hence went for the option most beneficial to metabo-GrimAge. To make the calibration plots, we considered 5-year mortality and placed participants in five equally sized groups based on their predicted 5-year mortality.

The results for the ‘traditional’ evaluation of biological age clocks, i.e. to what extent Δ is associated with all-cause mortality in a Cox PH model that also includes chronological age and sex, can be found in Table 1. It can be seen that the Δ of both metabo-AccelerAge-Gompertz and metabo-GrimAge is strongly associated with time to all-cause mortality.

**Table 1.** Hazard ratios for all-cause mortality associated with a standard unit increase in Δ in a Cox PH model adjusted for chronological age and sex, evaluated on the test set of the UKB data.

|  | HR (95% CI) | p-value |
| --- | --- | --- |
| $\Delta$: metabo-GrimAge | 1.61 (1.56–1.65) | < 2e-16 |
| $\Delta$: metabo-AccelerAge-Gompertz | 1.56 (1.52–1.60) | < 2e-16 |

Table 2 contains the results for one of our new proposed evaluation measures, the concordance. Both biological age predictors have a higher concordance than chronological age.

**Table 2.**
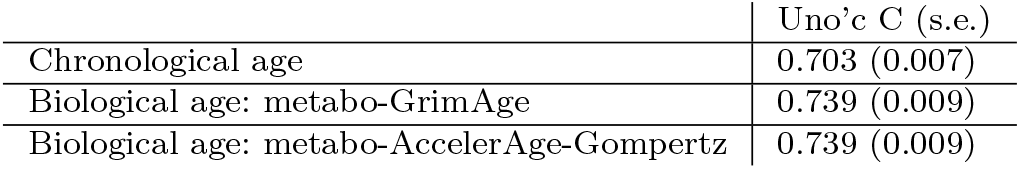
Concordance (Uno’s C) of the biological age predictions evaluated on the test set of the UKB data.

Calibration plots for metabo-AccelerAge-Gompertz and metabo-GrimAge can be found in Figure 9. AccelerAge-Gompertz is better calibrated: predicted all-cause mortality risk is closer to observed all-cause mortality risk. This result can be understood when plotting chronological age against predicted biological age, as done in Figures 10 and 11. The metabo-GrimAge predictions are—by design—centered around the line *C* = *B*. For this particular data set that does not make sense, since we know that the UKB population lives longer than the average UK population (Figure 8), so their biological ages should on average be somewhat lower than the line *C* = *B*. The predicted biological ages of our metabo-AccelerAge-Gompertz predictor are on average also lower than the chronological ages.

**Fig. 9.**
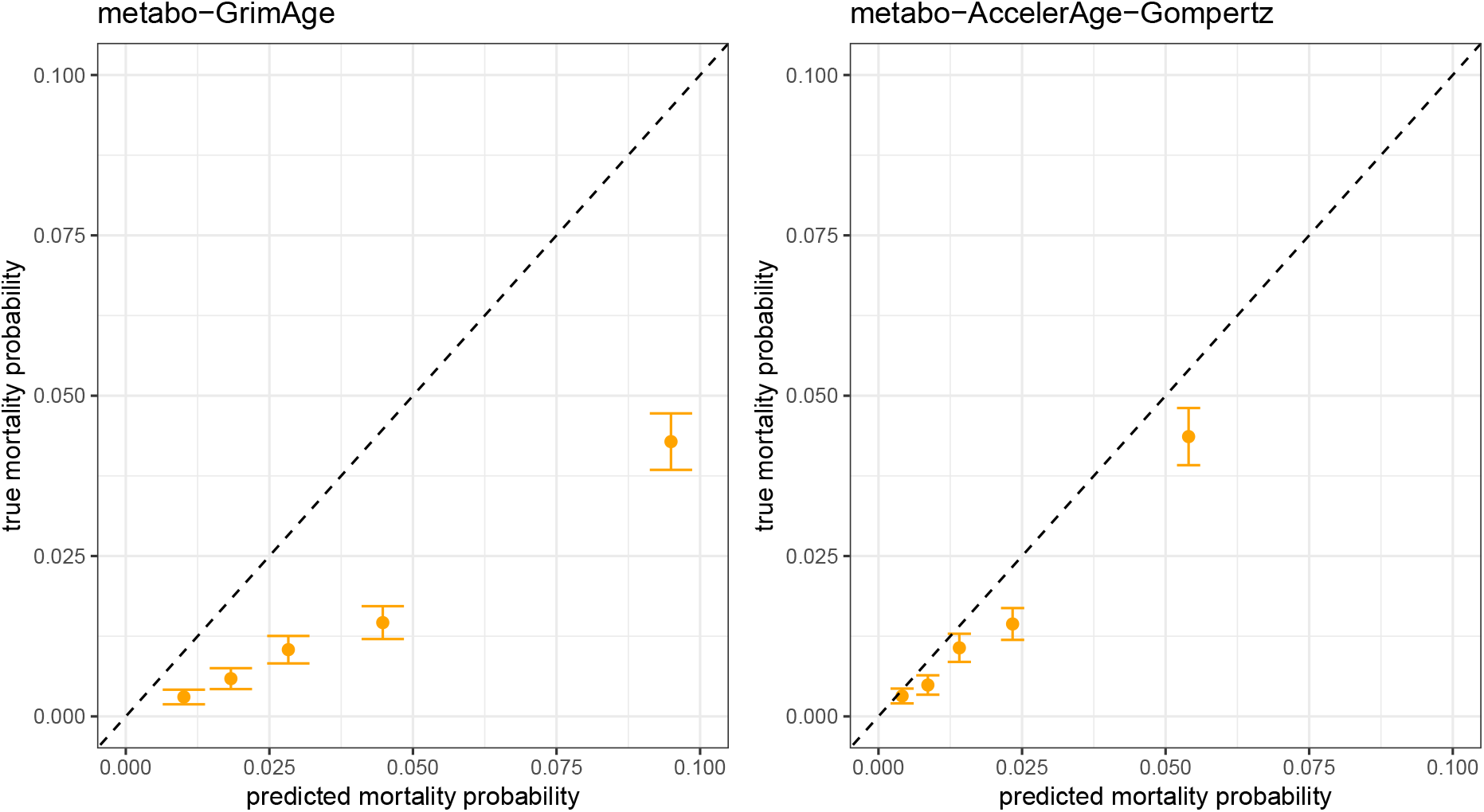
Calibration metabo-GrimAge and metabo-AccelerAge-Gompertz based on 5-year survival in the test set of the UKB data.

**Fig. 10.**
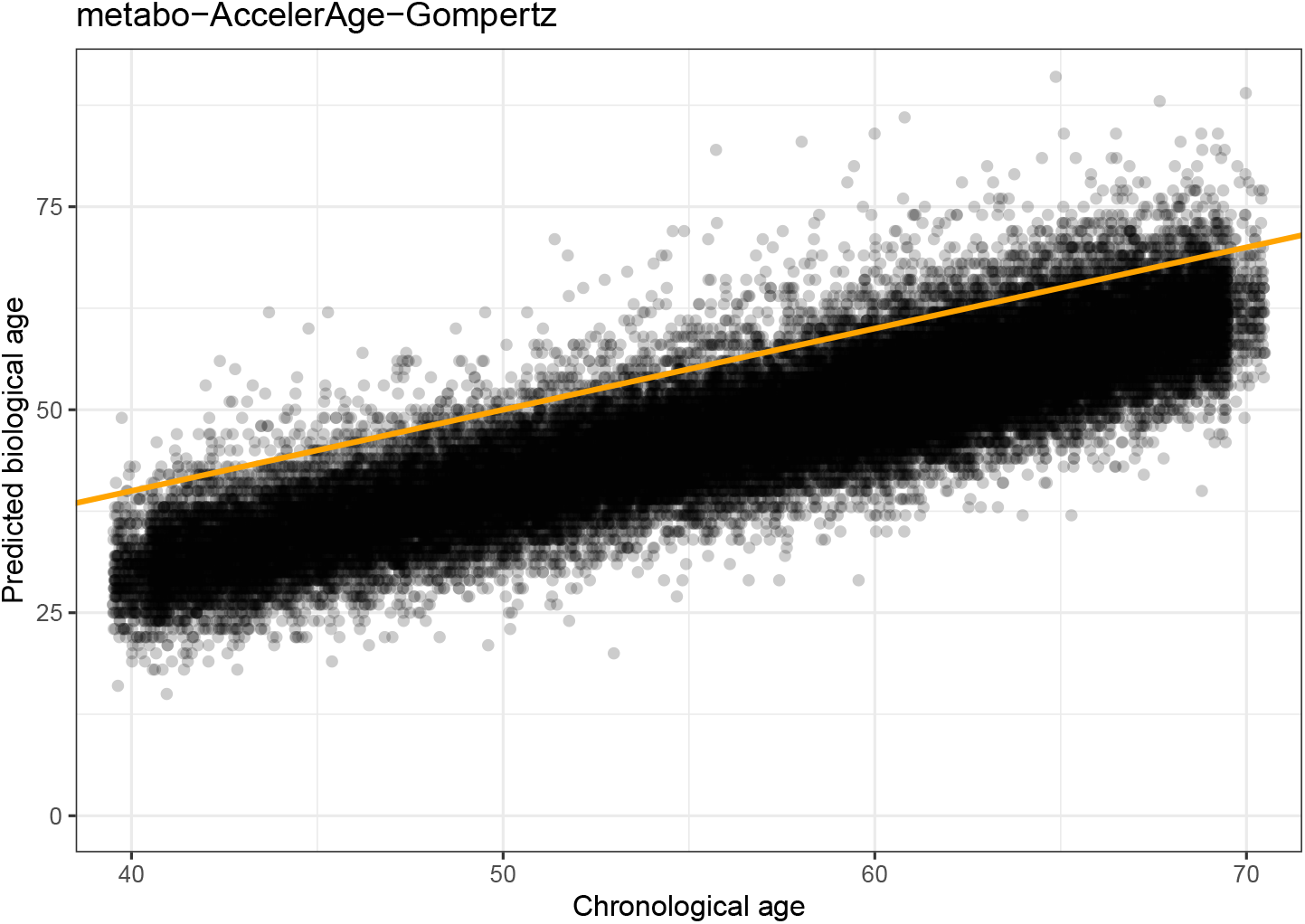
Chronological age plotted against predicted biological age in the test set of the UKB data for metabo-AccelerAge-Gompertz. The orange line has slope 1: it denotes where chronological age is equal to predicted biological age.

**Fig. 11.**
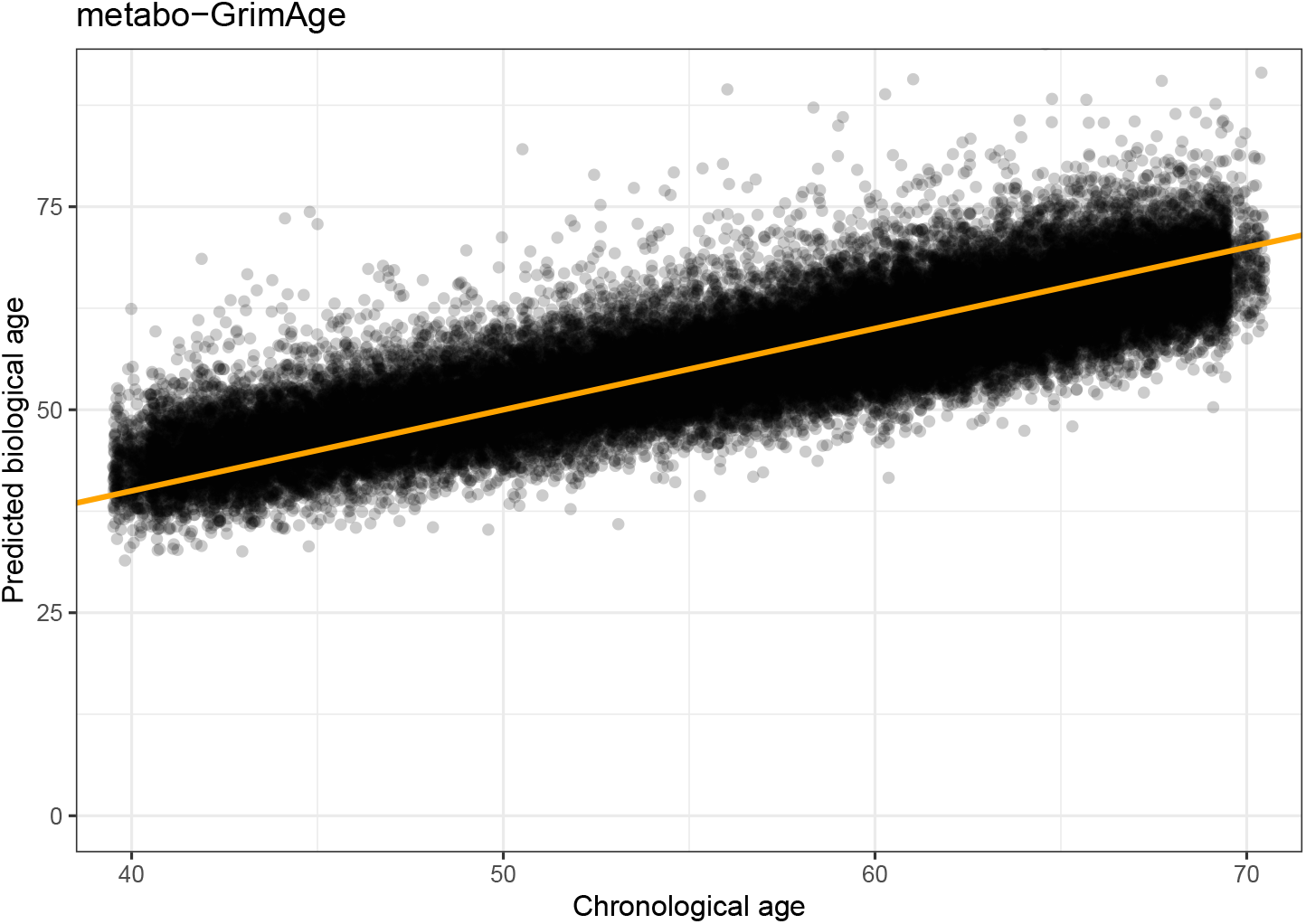
Chronological age plotted against predicted biological age in the test set of the UKB data for metabo-GrimAge. The orange line has slope 1: it denotes where chronological age is equal to predicted biological age.

We validated our findings that metabo-AccelerAge-Gompertz and metabo-GrimAge perform equally well on discrimination but that metabo-AccelerAge-Gompertz is better calibrated in an external data set, the Leiden Longevity Study (LLS). The LLS tracks long-lived Dutch siblings of Caucasian origin, their offspring and the partners of the offspring. Participants were recruited between March 2002 and May 2006. Registry-based follow-up data until November 2021 was available. We used data on the offspring and the partners (N = 2312). Participants who dropped out (N = 10) or had missing values on the 242 included metabolic variables (N = 46) were excluded, resulting in 1007 men and 1249 women with a mean age of 59.2 years (sd 6.7) at inclusion. Median follow-up time was 16.3 years (IQR: 15.3–17.1) and 313 deaths were observed.

Table 3 shows to what extent age acceleration Δ is associated with all-cause mortality in a Cox PH model that also includes chronological age and sex on the LLS data. The Δ-values of both biological age predictors are still significantly associated with mortality beyond chronological age. Nevertheless, the hazard ratios are much lower.

**Table 3.** Hazard ratios for all-cause mortality associated with a standard unit increase in Δ in a Cox PH model adjusted for chronological age and sex, evaluated on the LLS data.

|  | HR (95% CI) | p-value |
| --- | --- | --- |
| $\Delta$: metabo-GrimAge | 1.39 (1.23–1.57) | 1.58e-7 |
| $\Delta$: metabo-AccelerAge-Gompertz | 1.35 (1.21–1.52) | 1.82e-7 |

Table 4 contains the concordance of the biological age predictors on the LLS data. The two predictors still both discriminate slightly better than chronological age, but the difference is smaller.

**Table 4.**
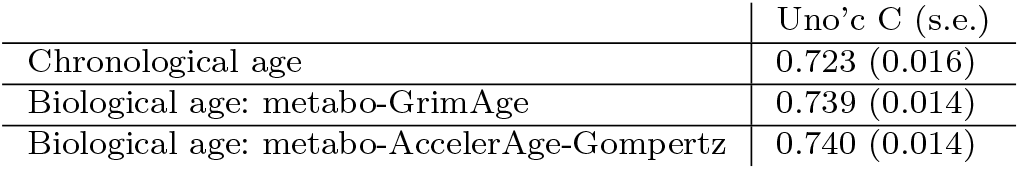
Concordance (Uno’s C) of the biological age predictions evaluated on the LLS data.

|  | Uno’s C (s.e.) |
| --- | --- |
| Chronological age | 0.723 (0.016) |
| Biological age: metabo-GrimAge | 0.739 (0.014) |
| Biological age: metabo-AccelerAge-Gompertz | 0.740 (0.014) |

Figure 12 contains the calibration plots for the two predictors on the LLS data. The metabo-AccelerAge-Gompertz predictor is still better calibrated than metabo-GrimAge, although the difference is smaller. In this case the biological age predictions of metabo-AccelerAge-Gompertz are slightly too low, especially for the group with the highest predicted mortality probability: the true event rates are slightly higher than the predicted probabilities.

**Fig. 12.**
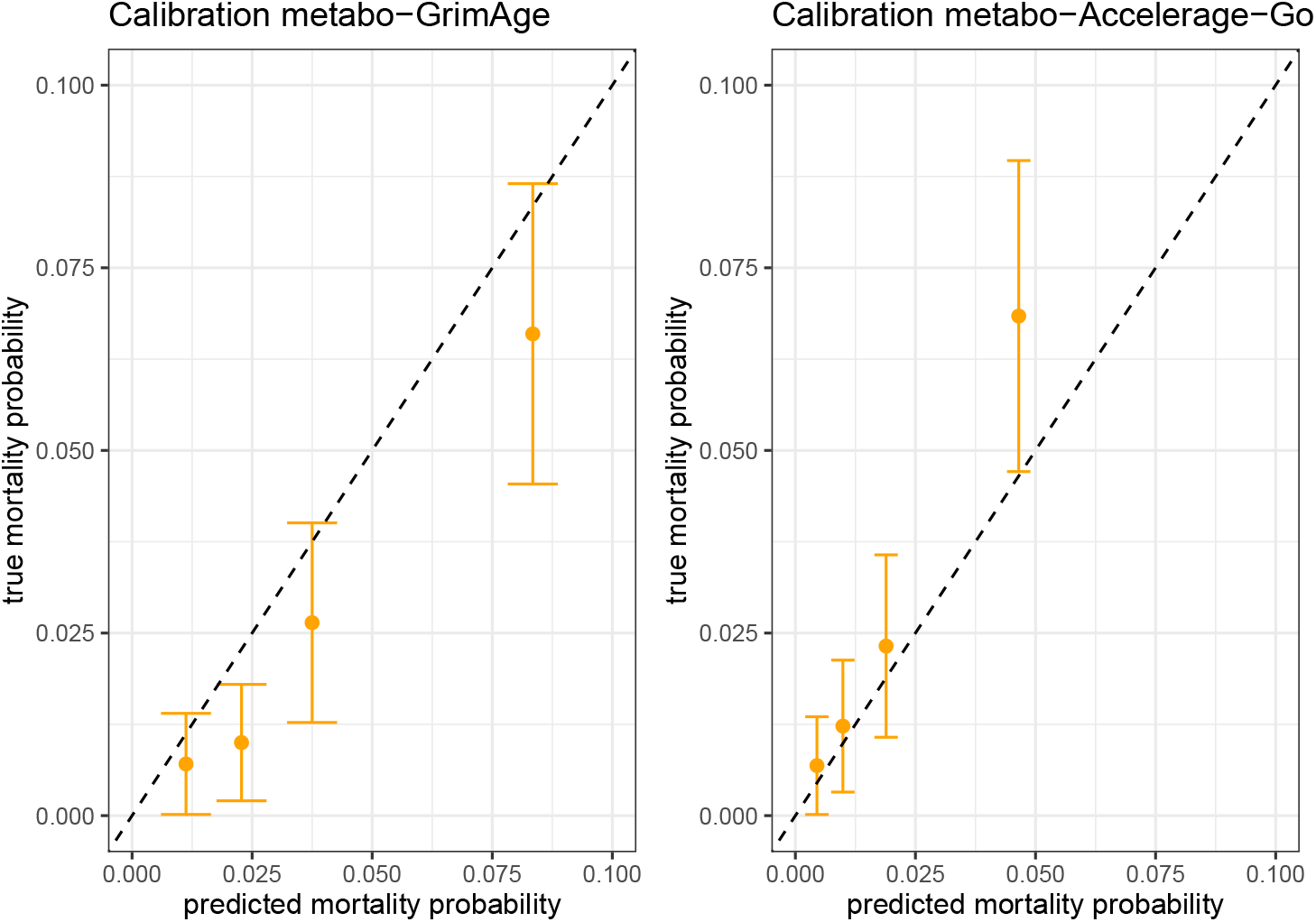
Calibration metabo-GrimAge and metabo-AccelerAge-Gompertz based on 5-year survival in the LLS data.

## 9 Discussion

In this paper we presented a new statistical framework for biological age prediction, AccelerAge, based on Accelerated Failure Time models. We proposed this new framework because current biological age predictors are not based on a formal operationalization of biological age, are not in line with the aging-as-a-clock idea and are difficult to properly evaluate. To ensure the AccelerAge framework is not just another ad hoc approach, we started by formally defining the concept of biological age via residual life, and subsequently basing AccelerAge on this operationalization. The discussion on what (biological) aging exactly entails is vivid and ongoing, but our proposed definition can serve as a starting point for further discussion. We explained why biological age predictors based on AFT models are in line with the ubiquitous clock metaphor in the aging field, while predictors based on Proportional Hazards models are not. Besides the more natural interpretation of biological age predictors based on AFT models, another advantage of AFT models is that they are robust to covariate omission: neglecting true covariates might lead to a distribution outside the parametric family considered, but it does not affect the regression-part of the model. This is not true for the PH model [33]. Another appealing aspect of the AccelerAge framework is that the AFT-model is fitted using chronological age as the timescale *t* instead of time-on-study. This is the preferred timescale in the context of cohort studies [34, 35]. Finally, in this paper we introduced two new evaluation measures into the context of biological age clocks, namely concordance and discrimination. This allows for a broader evaluation and validation of biological age predictors. AccelerAge predictions are directly interpretable on an age-scale, which means that the biological age prediction itself is meaningful, not just the age-independent part Δ.

Our UK Biobank application illustrates that metabo-AccelerAge-Gompertz is a worthy competitor to our GrimAge-implementation, metabo-GrimAge. Using the often-used evaluation measure of checking whether the age-independent part of a biological age prediction (Δ) is significantly associated with prospective mortality, the hazard ratio for metabo-AccelerAge-Gompertz was slightly lower than that for our GrimAge-implementation, but metabo-AccelerAge-Gompertz was better calibrated in both the UKB data and the LLS data, used as the external validation set. On concordance the two predictors scored similarly.

The real data application highlights the need for having more than one evaluation measure in biological age prediction. A true biological age predictor should be directly interpretable on an age-scale. Calibration plots can be used to assess the ability of a predictor to do so; checking whether Δ is significantly associated with time-to-mortality cannot. In addition, only relying on the association of Δ with agerelated outcomes might paint a too optimistic picture of the (clinical) usefulness of a biological age predictor. Although we found that both metabolite-based biological age clocks performed better than chronological age on discrimination, the differences with chronological age were small. One potential reason for this is that we only considered the metabolic variables as measured by the Nightingale NMR platform as predictor variables. Using more of the human metabolome, or using other sources of omics, might capture more of the aging process and hence further improve the discriminative ability of a predictor.

Our AccelerAge framework can be applied to any kind of time-to-event outcome and any type of predictor variables. Naturally, the conclusions resulting from the comparison with a GrimAge-based predictor might change if different predictor variables or outcomes are considered. We plan to extend the framework to also allow for regularization. Incorporating multiple predictor types, e.g. multiple omics, would be possible with a group-wise penalty term.

There are several limitations to our work. Our proposed operationalization is only based on lifespan, not healthspan. This can be considered a (too) narrow view of what it means to age. In addition, our operationalization requires that one has access to a lifetable of the reference population of interest. This might not always be the case, in particular not for outcomes other than mortality. In certain cases it might be feasible to simply construct a lifetable based on the training data set itself, but then the ability to detect whether the reference population differs from the sample is lost.

We illustrated the AccelerAge framework using a fully parametric AFT model, based on the Gompertz distribution. As it has long been known that the Gompertz distribution describes adult mortality well, this parametric approach sufficed for our real data application, in which time-to-mortality was our only outcome of interest. AFT models can also be fit using a semiparametric [47–50] or flexible parametric [51] approach. We initially included these approaches in our simulation study as well, but found that especially the fitting of flexible parametric AFT models can be inconsistent and suffer from convergence issues. Results for our simulation study where we also included a semiparametric and flexible parametric AccelerAge approach can be found in section 4 of the Supplementary Information. However, as developing robust flexible parametric AFT models is an area of active research [52], we believe the flexible parametric AccelerAge approach will soon become an appealing alternative to our fully parametric approach. It should however be noted that, in order to fit any AccelerAge model that is not fully parametric, the data must cover the whole of human lifespan to properly estimate the baseline survival curve or hazard function (which must be fully specified, because it needs to be integrated over). In the UK Biobank data, the oldest included participant was 85. Semiparametric prediction approaches therefore would have no data to estimate baseline survival or hazard at ages after 85.

In conclusion, our work represents a substantial advancement in the field of biological age research. By introducing AccelerAge, a new AFT-based statistical framework to predict biological age based on a solid operationalization of biological age, and incorporating previously underutilized evaluation measures, namely discrimination and concordance, we have laid a robust statistical foundation for the development and validation of biological age clocks.

## Statements and Declarations

### Supplementary Information

This manuscript has supplementary material available at [link to add]. R-code illustrating how to fit the AccelerAge framework is available in a public GitHub repository (https://github.com/marije-sluiskes/fitting-accelerage-framework-in-r).

### Funding

Marije Sluiskes was funded by the Netherlands Organization for Scientific Research, domain Health Research and Medical Sciences (09120012010052).

The Leiden Longevity Study has received funding from the European Union’s Seventh Framework Programme (FP7/2007-2011) under grant agreement number 259679. The LLS was further supported by a grant from the Innovation-Oriented Research Program on Genomics (SenterNovem IGE05007), the Centre for Medical Systems Biology, and the Netherlands Consortium for Healthy Ageing (grant 050-060-810), all in the framework of the Netherlands Genomics Initiative, Netherlands Organization for Scientific Research (NWO), by BBMRI-NL, a Research Infrastructure financed by the Dutch government (NWO 184.021.007 and 184.033.111) and the VOILA Consortium (ZonMw 457001001).

### Competing interests

The authors have no relevant financial or non-financial interests to disclose.

### Author contributions

Conception and design: Marije Sluiskes, Jelle Goeman, Hein Putter, Mar Rodŕıguez-Girondo. Provision of data: Marian Beekman, Eline Slagboom, Erik van den Akker. Writing - original draft preparation: Marije Sluiskes. Writing - review and editing: Jelle Goeman, Hein Putter, Mar Rodŕıguez-Girondo, Marian Beekman, Eline Slagboom. All authors read and approved the final manuscript.

### Ethics approval

The UK Biobank Study was approved by the North West Multi-Centre Research Ethics Committee (Reference Number 16/NW/0274). The Leiden Longevity Study was approved by the Medical Ethics Committee of the Leiden University Medical Center.

### Consent to participate

Informed consent was obtained from all individual participants included in the UK Biobank Study or the Leiden Longevity Study.

## Notes

https://github.com/marije-sluiskes/fitting-accelerage-framework-in-r

### Competing Interest Statement

The authors have declared no competing interest.

