## Supplementary Information for "The AccelerAge framework: A new statistical approach to predict biological age based on time-to-event data"

2023

#### 1 Gompertz distribution as a PH and as an AFT model

Although the Gompertz distribution is usually parameterized as a Proportional Hazards (PH) model, it can also be an Accelerated Failure Time (AFT) model (but, contrary to the Weibull distribution, not at the same time). However, it requires a different parameterization to see this (Broström [2021]), as will be illustrated below.

A regression model adheres to the proportional hazards assumption if it fulfills the assumption that

$$h(t) = h_0(t) \times \exp(\beta^T X), \tag{1}$$

where $h_0(t)$ is the baseline hazard. This is the proportional hazards assumption: the ratio of $h(t)/h_0(t)$ is the constant $\exp(\beta^T X)$.

A model adheres to the accelerated failure time assumption if it fulfills the assumption that

$$S(t|X) = S_0(t \times \exp(\beta^T X)), \tag{2}$$

where $S_0(t)$ is the baseline survival. The expression $\exp(\beta^T X)$ is also known as the 'acceleration factor' in the context of AFT models.

##### Gompertz as a PH model

The usual Gompertz parametrization is the rate parametrization, where the hazard is given by

$$h(t) = a \exp(bt). \tag{3}$$

Here, $a$ is generally called the rate parameter and $b$ the shape parameter. Now we have to choose how to insert the linear predictor and show that the resulting model adheres to assumption (1). For simplicity, assume $X$ is binary. Insert the linear predictor by reparameterizing $a = \exp(\beta^T X)$. Then the hazard ratio ($X = 1$ vs. $X = 0$) equals

$$HR = \frac{\exp(bt) \times \exp(\beta^T X)}{\exp(bt)} = \exp(\beta^T X), \tag{4}$$

such that the proportional hazards assumption of (1) is met.

##### Gompertz as an AFT model

To see that a Gompertz regression model can also be parameterized as an AFT model, rewrite the hazard as

$$h(t) = \frac{\tau}{\sigma} \exp(\frac{t}{\sigma}), \tag{5}$$

where $\sigma = \frac{1}{b}$ and $\tau = \frac{a}{b}$. Then the survival function can also be written in canonical form:

$$S(t) = \exp[-\tau(\exp(\frac{t}{\sigma} - 1)]. \tag{6}$$

The next step is again to insert the linear predictor somewhere and show that the resulting model adheres to 2. For simplicity, assume $X$ is binary. Take $\frac{1}{\sigma}$ as the linear predictor $\exp(\beta^T X)$. Then

$$S(t|X=1) = \exp[-\tau(\exp(t \times \exp(\beta^T X)) - 1)] \tag{7}$$

and

$$S(t|X=0) = \exp[-\tau(\exp(t-1)] = S_0(t). \tag{8}$$

Now note that

$$S_0(t \times \exp(\beta^T X)) = \exp[-\tau(\exp(t \times \exp(\beta^T X)) - 1)], \tag{9}$$

which is equal to expression (7). Hence, assumption (2) is satisfied. If the linear predictor would have been inserted in a different way, for example taking $\sigma$ as the linear predictor $\exp(\beta^T X)$, then the above equation would not have been equal to (7).

#### 2 GrimAge in detail

In both the simulation study as well as the real data application we fitted our own version of GrimAge, because we do not use DNAm data as predictor variables. Therefore we could not use the weights of the published GrimAge predictor, but we repeated the approach used to construct the original GrimAge predictor with different predictor variables. Here we describe our implementation and the differences with the approach to fit the original GrimAge predictor.

In the original publication [Lu et al. 2019], the GrimAge predictor is constructed following a two-step approach. In the first stage, DNAm-based surrogate biomarkers of smoking pack-years and a selection of plasma proteins are defined. In the second stage, together with chronological age and sex these surrogate biomarkers are included in an elastic net Cox Proportional Hazards model with time to all-cause mortality as outcome, after which the linear predictors of this model are transformed to an age-scale. The authors justify this two-step approach because the single-step approach (directly regressing CpG-sites on time to all-cause mortality) resulted in a less significant p-value. The transformation to an age-scale depends on the mean and standard deviation of chronological age in the dataset that the GrimAge predictor is fitted on.

Measurements of individual CpG-sites are known to be quite unreliable [Sugden et al. 2020]. A solution to this was recently suggested by Higgins-Chen et al. [2021], who used principal components as predictors: the first step of defining surrogate markers is hence replaced by the step of finding the principal components. This resulted in more reliable DNAm-based clocks and has the added benefit of creating a set of uncorrelated predictors. When fitting our version of GrimAge, metabo-GrimAge, on the UK Biobank data we hence followed this approach: we did not construct surrogate markers, but did a principle component analysis on our candidate markers of aging and included these principle components as the predictor variables in step 2. (In the simulation study, we only considered two independently generated predictor variables, so no dimension reduction step was required in the first place.) Our step 2 is exactly similar to step 2 in the original GrimAge model-fitting approach.

##### First stage

- Perform principal component analysis on the set of predictor variables.
- Include the first $i$ principal components that collectively explain at least 95% of the variance as predictor variables in step 2.

##### Second stage

- On the training data, fit a Cox PH model with follow-up time as the outcome variable (i.e. time-on-study as timescale $t$) and $C$, sex and the principal components from step 1 as the predictor variables.
- Obtain the linear predictors: $\hat{\beta}^T X_{train}$.
- Determine coefficients $a$ and $b$ such that the mean and standard deviation of the linearly transformed linear predictors $a + b(\beta^T X_{train})$ are the same as the mean and standard deviation of $C$ in the training data.
- Obtain linear predictors for the data set of interest (the test data, or some new data set).
- Linearly transform these linear predictors using the values for $a$ and $b$ as estimated earlier. This results in the GrimAge prediction:
$$GrimAge = a + b(\hat{\beta}^T X_{test}).$$
- To get $\hat{\Delta}$: regress GrimAge on $C$ and obtain the residuals.

#### 3 How to draw survival times

##### Proportional Hazards

Following Bender et al. [2005], the survival function of a Cox PH model is given by

$$S(t|X) = \exp[-H_0(t) \times \exp(\beta^T X)] \tag{10}$$

and hence the cumulative distribution function is given by

$$F(t|X) = 1 - S(t|X) = 1 - \exp[-H_0(t) \times \exp(\beta^T X)]. \tag{11}$$

Let $Y$ be a random variable with distribution function $F$. Then $U = F(Y) \sim U[0,1]$. Let $T$ be the survival times from the Cox model. It follows that

$$F(T|X) = 1 - \exp[-H_0(T) \times \exp(\beta^T X)] = U. \tag{12}$$

If $U \sim U[0,1]$ then $(1-U) \sim U[0,1]$. Hence,

$$U = \exp[-H_0(T) \times \exp(\beta^T X)]. \tag{13}$$

Finally, solve (13) for $T$:

$$t_i = H_0^{-1}\left[\frac{-\log(U)}{\exp(\beta^T X_i)}\right]. \tag{14}$$

##### Accelerated Failure Time

The survival function of an AFT model is given by

$$S(t|X) = H_0[t \times \exp(\beta^T X)], \tag{15}$$

and hence the cumulative distribution function is given by

$$F(t|X) = 1 - S(t|X) = 1 - H_0[t \times \exp(\beta^T X)]. \tag{16}$$

Let $Y$ be a random variable with distribution function $F$. Then $U = F(Y) \sim U[0,1]$. Let $T$ be the survival times from the AFT model. It follows that

$$F(T|X) = 1 - H_0[T \times \exp(\beta^T X)] = U. \tag{17}$$

If $U \sim U[0,1]$ then $(1-U) \sim U[0,1]$. Hence,

$$U = H_0[T \times \exp(\beta^T X)]. \tag{18}$$

Finally, solve (18) for $T$:

$$t_i = \frac{H_0^{-1}[-\log(U)]}{\exp(\beta^T X_i)}. \tag{19}$$

Table 1: Number of simulation runs (out of $n_{sim} = 200$) for which the flexible parametric AFT model (AccelerAge-flexpar) did not properly converge and which were subsequently excluded from the results.

| $n_{obs}$ | Gompertz PH | Gompertz AFT | Weibull |
|---|---|---|---|
| 500 | 9 | 15 | 20 |
| 2,500 | 15 | 30 | 47 |
| 5,000 | 27 | 69 | 45 |
| 7,500 | 34 | 97 | 51 |
| 10,000 | 51 | 135 | 49 |

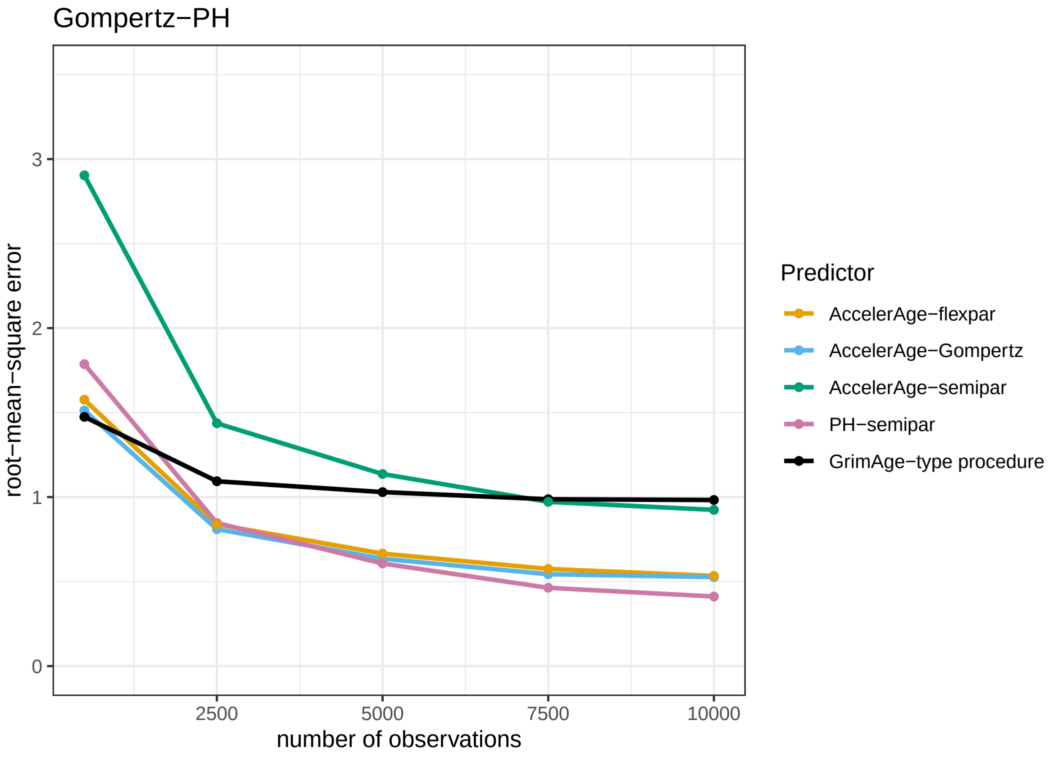


Figure 1: Performance of the five different biological age predictors in terms of the root-mean-square error under the Gompertz-PH data-generating mechanism. Results are reported for data sets of varying sizes ($n_{obs} = 500$, 2500, 5000, 7500 and 10,000) as the average root-mean-square error over a simulation sample size of $n_{sim} = 200$.

### 4 Simulation study results semiparametric and flexible parametric AFT

This section of the supplementary material contains the results of a simulation study that includes two additional predictors, based on a semiparametric and flexible parametric AccelerAge approach. All other details of the simulation study are exactly as described in the main manuscript.

Results can be found in Figures 1, 2 and 3 As mentioned, the flexible parametric AFT model is numerically delicate to fit and often had troubles to converge. We excluded simulation runs for which this occurred from the results. Table 1 contain the number of simulation runs (out of $n_{sim} = 200$ for which this was the case. Results are hence based on fewer simulation runs, but when comparing the figures in this section with those in the main manuscript, it can be seen that the performance of the three methods that are included in both is almost identical. Especially the Gompertz AFT model proved to be problematic. This model is numerically delicate to fit.

To fit the semiparametric AFT, we use the approach of Stute [1993], described in more detail in the next section of this document. The flexible parametric AFT models were fitted using the function `aft()` from the R-package rstpm2 [Liu et al. 2018] with `df = 3`.

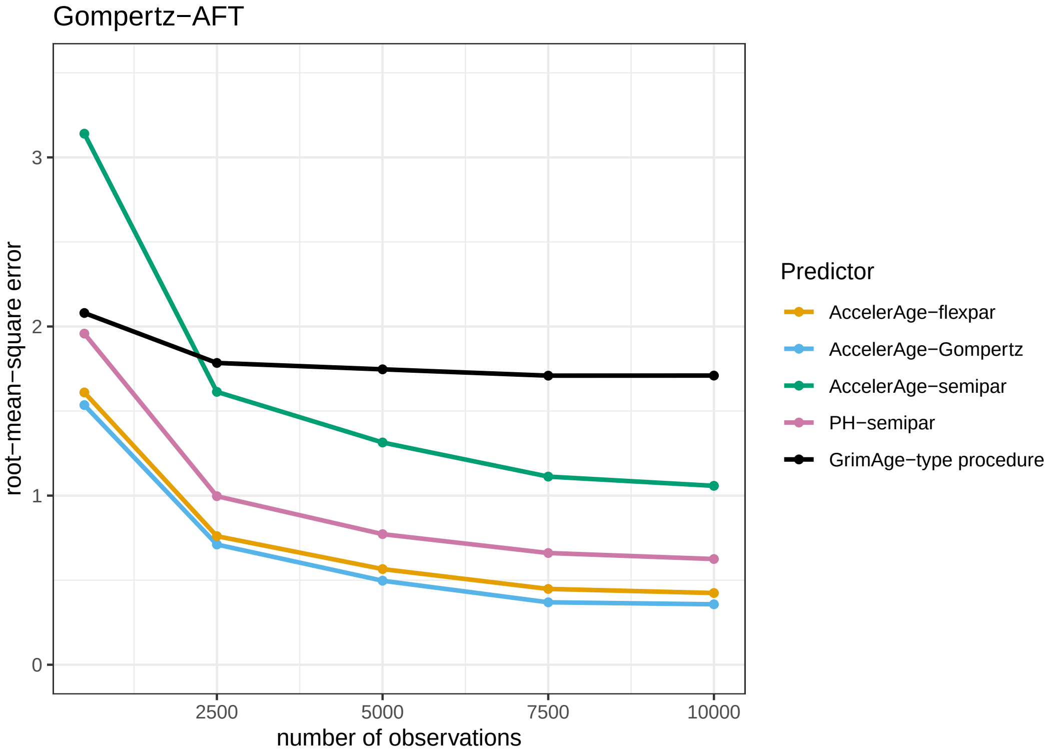


Figure 2: Performance of the five different biological age predictors in terms of the root-mean-square error under the Gompertz-AFT data-generating mechanism. Results are reported for data sets of varying sizes ($n_{obs} = 500$, 2500, 5000, 7500 and 10,000) as the average root-mean-square error over a simulation sample size of $n_{sim} = 200$.

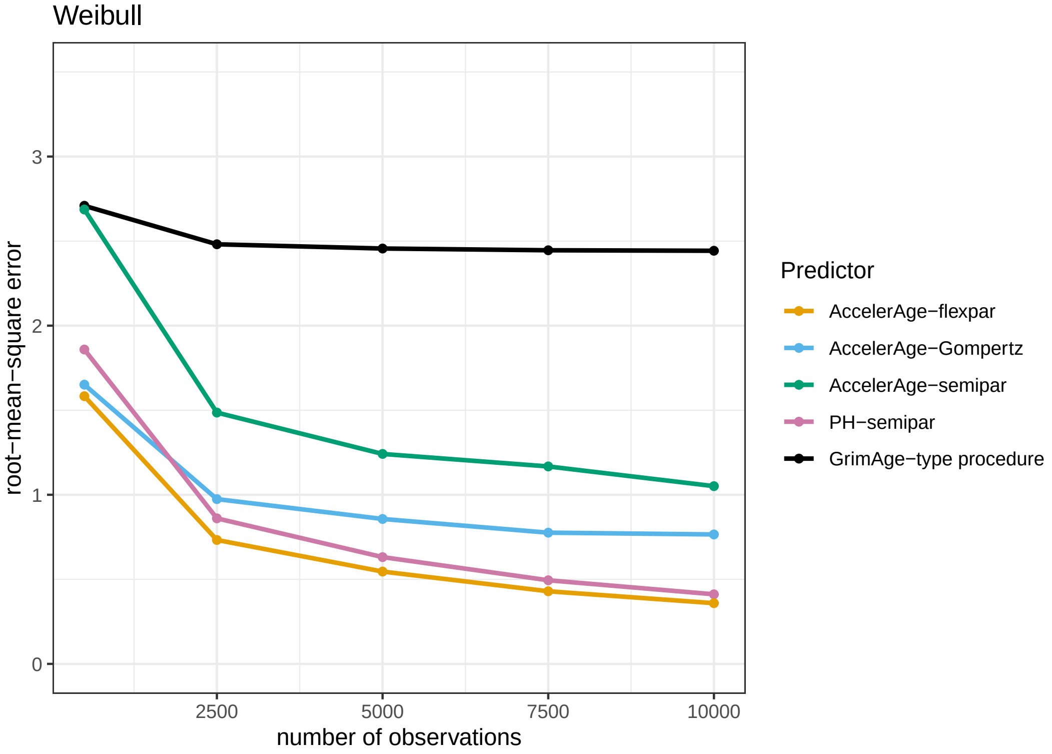


Figure 3: Performance of the five different biological age predictors in terms of the root-mean-square error under the Weibull data-generating mechanism. Results are reported for data sets of varying sizes ($n_{obs} = 500$, 2500, 5000, 7500 and 10,000) as the average root-mean-square error over a simulation sample size of $n_{sim} = 200$.

### 5 The semiparametric AFT

The semiparametric AFT model does not assume an underlying parametric baseline hazard so this needs to be estimated. We use the weighted least squares method, based on Kaplan-Meier weights, as suggested by Stute [1993]. Estimates for the coefficients are found via:

$$\hat{\beta} = \arg\min \sum_{i=1}^{n} W_i^L [\log(T_{(i)}) - \beta^T X_{(i)}]^2, \tag{20}$$

where $\log(T_{(i)})$ is the $i^{th}$ ordered value of the observed response $\log(T)$ and $X_{(i)}$ is the corresponding covariate. If there is no delayed entry, $W$ contains the Kaplan-Meier weights (the successive increments of the Kaplan-Meier estimator) of $T_{(i)}$. If there is, $W$ is adjusted to:

$$W_i^L = W_i / \sum_{j=1}^{n} I(L_j < T_i < T_j), \tag{21}$$

where $L_j$ is the left truncation time of individual $j$. In other words, the risk set is adjusted. Note that since the Kaplan-Meier estimator does not change when an individual is censored, $W_i/W_i^L$ is zero for each censored individual.

Once estimates for $\beta$ are obtained, an estimate for the baseline survival is next. First of all, remember that:

$$\log(T_{(i)}) - \beta^T X_{(i)} = \epsilon_{(i)} \tag{22}$$

and that

$$\hat{F}_0(u) = \hat{P}(\omega \leq u) = \sum_{i=1}^{n} W_i I(\exp(\log(T_{(i)}) - \hat{\beta}^T X_{(i)}) \leq u) = \sum_{i=1}^{n} W_i I(\hat{\epsilon}_{(i)} \leq u). \tag{23}$$

Now, $\hat{S}_0(u)$ can be obtained by $1 - \hat{F}_0(u)$. To get this to the timescale of interest, remember that $\log(T_{(i)}) = \beta^T X_{(i)} + \epsilon_{(i)}$, so $T_{(i)} = \exp(\beta^T X_{(i)} + \epsilon_{(i)}) = \exp(\beta^T X_{(i)}) \times \exp(\epsilon_{(i)})$. For the baseline, set all covariates to 0. The intercept then remains, hence $T_{0,(i)} = \exp(\beta_0) \times \exp(\epsilon_{(i)})$.
